# The microbial metabolite *p*-Cresol induces autistic-like behaviors in mice by remodeling the gut microbiota

**DOI:** 10.1101/2020.05.18.101147

**Authors:** P. Bermudez-Martin, J. A. J. Becker, N. Caramello, S. P. Fernandez, R. Costa-Campos, J. Canaguier, S. Barbosa, L. Martinez-Gili, A. Myridakis, M.-E. Dumas, A. Bruneau, C. Cherbuy, P. Langella, J. Callebert, J.-M. Launay, J. Chabry, J. Barik, J. Le Merrer, N. Glaichenhaus, L. Davidovic

**Affiliations:** Université Côte d’Azur, Centre National de la Recherche Scientifique, Institut de Pharmacologie Moléculaire et Cellulaire, Valbonne, France; Université François Rabelais, INRAE, CNRS, INSERM, Physiologie de la Reproduction et des Comportements, IFCE, 37380, Nouzilly, France; Imperial College London, Division of Systems Medicine, Department of Metabolism, Digestion and Reproduction, Faculty of Medicine, London, SW7 2AZ, United Kingdom; Imperial College London, Genomic and Environmental Medicine, National Heart & Lung Institute, Faculty of Medicine, London, SW3 6KY, United Kingdom; European Genomic Institute for Diabetes, CNRS UMR 8199, INSERM UMR 1283, Institut Pasteur de Lille, Lille University Hospital, University of Lille, 59045 Lille, France; McGill University and Genome Quebec Innovation Centre, 740 Doctor Penfield Avenue, Montréal, QC, H3A 0G1, Canada; Université Paris-Saclay, AgroParisTech, INRAE, Institut Micalis, Jouy-en-Josas, France; INSERM, UMR-S 942, Department of Biochemistry, Lariboisière Hospital, Paris, France; Centre for Biological Resources, BB-0033-00064, Lariboisière Hospital, Paris, France

**Keywords:** microbiota, autism, behavior, reward system, metabolite, *p-*Cresol, 4-Cresol

## Abstract

**Background:** Autism Spectrum Disorders (ASD) are associated with dysregulation of the microbiota-gut-brain axis resulting in changes in microbiota composition as well as fecal, serum and urine levels of microbial metabolites. Yet, a causal relationship between dysregulation of the microbiota-gut-brain axis and ASD remains to be demonstrated. Here, we hypothesized that the microbial metabolite *p*-Cresol, which is more abundant in ASD patients compared to neurotypical individuals, could induce ASD-like behavior in mice.

**Results:** Mice exposed to *p*-Cresol for 4 weeks in drinking water presented social behavior deficits, stereotypies, and perseverative behaviors, but no changes in anxiety, locomotion, or cognition. Abnormal social behavior induced by *p*-Cresol was associated with decreased activity of central dopamine neurons involved in the social reward circuit. Further, *p*-Cresol induced changes in microbiota composition and social behavior deficits could be transferred from *p*-Cresol-treated mice to control mice by fecal microbiota transplantation (FMT). We also showed that mice transplanted with the microbiota of *p*-Cresol-treated mice exhibited increased fecal *p-*Cresol levels compared to mice transplanted with the microbiota of control mice and identified possible *p*-Cresol bacterial producers. Lastly, the microbiota of control mice rescued social interactions, dopamine neurons excitability and fecal *p*-Cresol levels when transplanted to *p-*Cresol-treated mice.

**Conclusions:** The microbial metabolite *p-*Cresol induces ASD core behavioral symptoms in mice via a gut microbiota-dependent mechanism. Our study paves the way for therapeutic interventions targeting the microbiota to treat patients with ASD.

## INTRODUCTION

ASD are frequent (1:68) neurodevelopmental disorders resulting from interactions between genetic predisposition and environmental risks [1]. While 10-25% of ASD cases are explained by mutations in specific genetic *loci*, twin studies have revealed that genetic and environmental factors share equal influence on ASD risk [2]. The identification of environmental factors contributing to ASD is therefore critical to better understand their etiology. ASD core symptoms encompass social interaction and communication deficits, perseverative/stereotyped behaviors, restricted interests and abnormal sensory processing [3]. Also, ASD often co-occur with anxiety disorder, hyperactivity and intellectual disability [3]. Several evidences support an involvement of the gut-microbiota-brain axis in ASD [1]. First, ASD is associated with gastrointestinal (GI) dysfunction and increased intestinal permeability [1]. Second, children with ASD and concurrent GI symptoms exhibit more pronounced social impairments, sensory over-responsivity and anxiety compared to ASD peers without GI symptoms [4–6]. Third, ASD patients exhibit gut microbiota dysbiosis. Indeed, bacterial β-diversity changes associated with increased and decreased abundances of the genera *Clostridioides* and *Bifidobacterium* respectively have been reported in three independent meta-analyses [7–9]. In a pilot study, fecal microbiota transfer (FMT) from healthy individuals to ASD patients durably alleviated both GI symptoms and ASD core symptoms [10]. Fourth, dysbiosis in ASD patients is associated with altered levels of urinary, plasmatic or fecal microbial metabolites such as methylamines, indoles and tyrosine-derived metabolites [11–18].

Altered microbiota composition [19–23] and abnormal levels of microbial metabolites [19, 22, 23] have also been observed in rodent models of ASD: the maternal immune activation (MIA), the diet-induced obesity (DIO) and the valproate environmental models, the BTBR idiopathic model, and the *Shank3b-*KO genetic model. Further, changes in microbiota composition induced by FMT or probiotic treatment alleviated behavioral alterations in several of these ASD models [19–21]. Finally, mice born from mothers transplanted with feces from ASD patients exhibited social behavior deficits, changes in microbiota composition and abnormal patterns of microbial metabolites [24]. Several microbial metabolites were shown to induce behavioral changes when administered to rodents. Treatment with the SCFA propionate induced social interaction deficits, stereotypies, cognitive deficits and anxiety in rats [16]. The tyrosine degradation product 4-ethylphenylsulfate (4-EPS) and indole induced anxiety in mice [1, 19, 25]. Altogether, these data suggested that dysbiosis could contribute to ASD core and associated symptoms via the production of microbial metabolites.

Among the microbial metabolites linked to ASD, the small aromatic metabolite *p*-Cresol (*para-* Cresol, 4-Cresol, 4-methylphenol) was consistently found at increased levels in the urine and feces of ASD patients [13, 14, 17, 18, 26]. Also, *p*-Cresol urinary levels correlated with the severity of ASD behavioral alterations [13, 14]. Although environmental exposure to *p*-Cresol is relatively common and occurs through the skin, as well as the GI and respiratory systems [27], the largest and most widespread source of this compound results from tyrosine degradation by the intestinal microbiota in the colon. At least 55 bacterial strains present in the human microbiota can produce *p-*Cresol. These strains are phylogenetically divergent spreading across the *Bifidobacteriaceae, Enterobacteriaceae, Coriobacteriaceae, Bacteroidaceae, Fusobacteriaceae*, *Lactobacillaceae*, and *Clostridiaceae* families [28]. It was also shown that *p*-Cresol conferred a selective growth advantage to *Clostridioides difficile* [29], possibly contributing to the overabundance of members of the *Clostridioides* genus in ASD patients [7–9]. However, to date, the causal relationship between microbiota dysbiosis, *p*-Cresol production and ASD symptoms has not been demonstrated. Here, we show that mice chronically exposed to *p*-Cresol presented social behavior deficits, stereotyped/perseverative behaviors, changes in microbiota composition, and elevated *p-*Cresol. Further fecal microbiota transplantation (FMT) experiments revealed that *p-*Cresol induced social deficits by a microbiota-dependent mechanism.

## RESULTS

### Mice exposed to *p-*Cresol exhibit ASD-like behaviors that persist after treatment discontinuation

To mimic exposure to *p*-Cresol through the GI tract, we treated C57BL/6J male mice with *p*-Cresol in drinking water starting at 4.5 weeks of age (Fig. 1A, Fig. S1A). A treatment for 4 weeks with *p*-Cresol did not induce changes in body weight, drink or food intake (Fig. S1B-D). Compared to untreated control animals, *p*-Cresol-treated mice exhibited increased levels of *p*-Cresol in urine and feces (Fig. S1E, F), but not in serum (Fig. S1G).

**Fig. 1.**
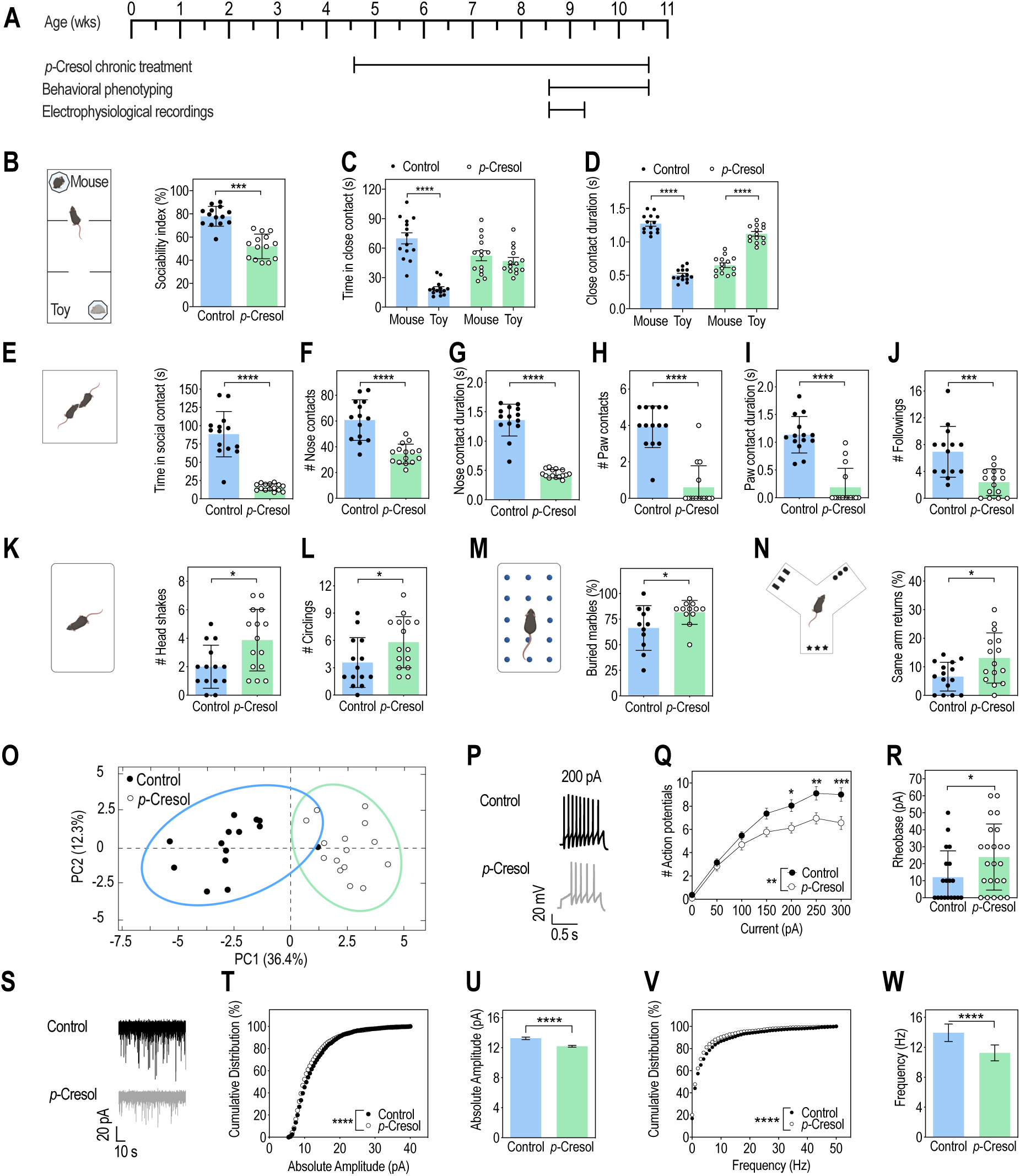
*p-*Cresol treatment induces autistic-like behaviors and alters VTA dopamine neurons excitability. (A) Timeline of the experiments. (B-D) Three-chamber test (n=14/group): (B) Sociability index (ratio of the time spent in close contact with the mouse interactor normalized to the sum of the time in close contact with both mouse interactor and toy mouse); Mann-Whitney U-test: ***p<0.001. (C) Time spent in close contact with the mouse interactor or the toy mouse; 2-way ANOVA: p(Treatment)=0.2601, p(Preference)=p<0.0001, p(Treatment x Mouse-Toy)=p<0.0001; Šidák’s post hoc tests for treatment effect: ****p<0.0001. (D) Mean duration of each close contact with the mouse interactor or the toy mouse; 2-way ANOVA: p(Treatment)=0.9416, p(Preference)=p<0.0001, p(Treatment x Mouse-Toy)=p<0.0001, Šidák’s post hoc tests for treatment effect: ****p<0.0001. (E-J) Dyadic social interaction test (n=14 Control, n=15 *p*-Cresol): (E) total time spent, (F) number of events and (G) mean duration of nose contacts, (H) number of events and (I) mean duration of paw contacts, (J) number of following events; Mann-Whitney U-test: ****p<0.0001, ***p<0.001. (K-L) Motor stereotypies (n=14 Control, n=15 *p*-Cresol): (K) number of head shakes and (L) circling events; Mann-Whitney U-test: *p<0.05. (M) Marble burying test (n=11 Control, n=12 *p-*Cresol): percentage of buried marbles; Mann-Whitney U-test: *p<0.05. (N) Y-maze spontaneous alternations (n=15/group): % of same arm returns; Mann-Whitney U-test: *p<0.05. (O) PCA plot of behavioral scores recorded in the dyadic social interaction and direct monitoring of motor stereotypies (Figure 1 E-L and Supplementary Figure 2H-L); ellipses of the 95% confidence intervals are indicated for each group (n=14 Control, n=15 *p*-Cresol); PC, principal component. (P-W) Electrophysiological recordings of VTA dopamine neurons activity: (P) Representative traces of dopamine neuron activity in patch-clamp experiments after a 200 pA current injection. (Q) Number of action potentials evoked by current injection in current-clamp experiments; n=5 animals/group, 19 cells recorded for Control, 23 cells recorded for *p*-Cresol; 2-way ANOVA: p(Treatment)=0.0066, p(Current)<0.0001, p(Treatment x Current)=0.0006; Šidák’s post hoc tests for treatment effect, *p<0.05,**p<0.01, ***p<0.001. (R) Rheobase, minimal electric current required to evoke an action potential; n=5 animals/group, 19 cells recorded for Control, 23 cells recorded for *p*-Cresol; Mann-Whitney U-test: *p<0.05. (S) Representative traces of spontaneous miniature excitatory postsynaptic currents (sEPSC) (T) Cumulative frequencies of sEPSC absolute amplitude; n=5 animals/group, >2500 events/group; Kolmogorov-Smirnov test: ****p<0.0001. (U) Mean absolute amplitude of sEPSC; Mann-Whitney U-test: ****p<0.0001. (V) Cumulative frequencies of sEPSC frequency; n=5 animals/group, >2500 events/group; Kolmogorov-Smirnov test: ****p<0.0001. (W) Mean frequency of sEPSC; Mann-Whitney U-test: ****p<0.0001. (B-N, R) Data are shown as means ± SD. (Q, U, W) Data are shown as means ± SEM.

*p-*Cresol-treated mice were subjected to behavioral phenotyping (Fig. 1A) for social interactions and repetitive/perseverative behaviors (as proxies for ASD core symptoms) (Fig. 1B-N, Fig. S2A-L) as well as anxiety, hyperactivity and cognitive deficits (as proxies of ASD comorbidities) (Fig. S2M-U). In the 3-chamber test (Fig. S2A-G, Fig. 1B-D), *p-*Cresol-treated mice presented reduced sociability (Fig. 1B) and no preference for the mouse interactor towards the toy mouse (Fig. 1C) compared to control mice. Although the number of close contacts with the mouse interactor was higher than with the toy mouse (Fig. S2G), their mean duration was reduced (Fig. 1D). During dyadic social interactions, *p*-Cresol-treated mice spent less time in social contact compared to control mice (Fig. 1E). The time, number and mean duration of both nose and paw contacts as well as the number of followings were also reduced (Fig. 1F-J, Fig. S2H, I) demonstrating that *p*-Cresol exposure impaired social interactions.

As for repetitive/perseverative behaviors, while *p*-Cresol-treated mice exhibited similar numbers of rearing episodes and time spent in digging and self-grooming compared to control mice (Fig. S2J-L), they displayed more frequent head shakes and circling events (Fig. 1K, L). An increase in stereotyped behaviors in *p*-Cresol-treated mice was confirmed in the marble burying test (Fig. 1M). *p*-Cresol-treated mice also exhibited increased frequency of perseverative same arm returns in the Y-maze spontaneous alternation task (Fig. 1N). Finally, *p*-Cresol-treated mice were clearly separated from control mice along the first principal component (PC1) axis in a PCA analysis of scores recorded in dyadic social interaction and direct monitoring of stereotypies (Fig. 1O).

As for other behaviors, *p-*Cresol-treated and control mice displayed similar nocturnal and diurnal locomotor activity as assessed in actimetry chambers, and travelled the same distance in the open-field (Fig. S2M-O), showing that *p*-Cresol exposure did not induce hyperactivity. Also, the number of entries and time spent in the open-field center, the latency to feed in the novelty-suppressed feeding test and the percentage of time spent in the open arm of the zero-maze were not impacted (Fig. S2P-S), suggesting that *p*-Cresol exposure did not induce anxiety. Finally, *p*-Cresol-treated mice explored objects similarly to control mice and displayed a similar recognition index for the novel object in the novel object recognition task, indicating that their cognitive ability were preserved (Fig. S2T, U).

We then investigated whether *p*-Cresol-induced behaviors were abolished when the treatment was discontinued (Fig. S3A). After a 4-week washout, the effect of *p*-Cresol on social interactions and stereotyped/perseverative behaviors persisted and were of similar magnitude (Fig. S3B-O). PCA analysis of social behavior and stereotypies scores revealed that control and *p-*Cresol-treated mice were clearly separated along the PC1 axis, both pre- and post-washout (Fig. S3P). Altogether, these results suggested that *p*-Cresol selectively induced ASD core symptoms which persisted long time after *p-*Cresol exposure.

### *p*-Cresol impairs VTA dopamine neurons excitability

We then investigated the impact of *p*-Cresol exposure on electrophysiological properties of dopamine neurons in the VTA, which are part of a “socially engaged reward circuit” [30]. Altered VTA connectivity and impaired VTA dopamine neurons excitability were observed in both ASD patients [31] and mouse models of ASD [20, 21, 32–35]. We used whole-cell patch-clamp to record VTA dopamine neurons in acute brain slices from control and *p*-Cresol-treated mice (Fig. 1P-W). First, *p*-Cresol-treated mice displayed reduced excitability of VTA dopamine neurons, with a reduction in the number of evoked action potentials (Fig. 1P-R). Second, both the amplitude and frequencies of miniature spontaneous excitatory postsynaptic currents (sEPSC) were reduced in VTA dopamine neurons of *p*-Cresol-treated animals compared to controls (Fig. 1S-W). Therefore, *p*-Cresol treatment resulted in decreased activity of VTA dopamine neurons.

### *p*-Cresol impacts the gut microbial ecology

Because *p-*Cresol exposure affects bacterial diversity [29], we analyzed the bacterial composition of the fecal microbiota in *p-*Cresol-treated and control mice using 16S ribosomal RNA (rRNA) sequencing. There was no difference in bacterial richness or evenness between the two groups, as assessed by number of amplicon sequence variant (ASV) and Shannon’s and Pielou’s indexes (Fig. 2A-C). In contrast, β-diversity analysis based on Aitchison’s distance revealed significant divergences in microbial composition between *p-*Cresol-treated and control mice (Fig. 2D). We then sought to identify discriminant features (ASV and bacterial taxa up to the phylum level) using the Analysis of Composition of Microbiomes (ANCOM) method [36] that considers the compositional nature of 16S datasets and exhibits good control of false discovery rate [37]. We found that 19 ASV, 4 species and 2 genera were discriminant (|centered-log ratio (CLR)|>0.2; W=0.7) with the largest effect size (|CLR|>0.5) being observed for ASV related to the *Bacteroidales, Clostridiales* and *Burkholderiales* orders (Fig. 2E, Tab. S1). We identified 15 taxonomic features increased in *p*-Cresol-treated mice (10 ASV, 3 species, 2 genera). In particular, *p*-Cresol increased the counts of ASV affiliated to one *Anaerobium* sp., *Turicimonas muris* and several *Muribaculaceae*: *Duncaniella dubosii*, one *Barnesiella* sp., one *Muribaculaceae bacterium,* one *Muribaculum* species (Fig. 2E,F, Tab. S1). We also identified 10 taxonomic features (9 ASV, 1 species), all affiliated to the *Clostridiales* order, that were depleted in *p*-Cresol-treated mice (Fig. 2E, Tab. S1). In particular, decreased counts of ASV affiliated to one *Eisenbergiella* sp., *Lacrimispora saccharolytica*, one *Clostridiaceae* bacterium, *Ruthenibacterium lactatiformans*, and one *Anaerotignum* sp. were observed in *p*-Cresol-treated mice (Fig. 2E, G, Tab. S1). Thus, *p-*Cresol-induced changes in microbial composition were mainly observed at the ASV level and scarcely at the species or genus level suggesting that *p-*Cresol induced selective and not broad taxonomic changes in microbial composition. To model how microbiota composition (assessed by ASV pseudocounts) predicted social abilities (Fig. 1E) and stereotypies (Fig. 1K, L), we then used a Random Forest (RF) regressor blind to the experimental groups. We found that 15 and 13 ASV were the strongest drivers of social abilities (Fig. 2H), and stereotypies (Fig. 2I) respectively. Of note, selected ASV explained social behavior scores, with up to 12% of the score explained by *Duncaniella dubosii* (Fig. 2H), while they contributed less to explain stereotypies with one *Clostridiaceae* bacterium explaining 5% of the score (Fig. 2I). These data reinforce the link between microbial composition and ASD-like core symptoms, and in particular social behavior deficits.

**Fig. 2.**
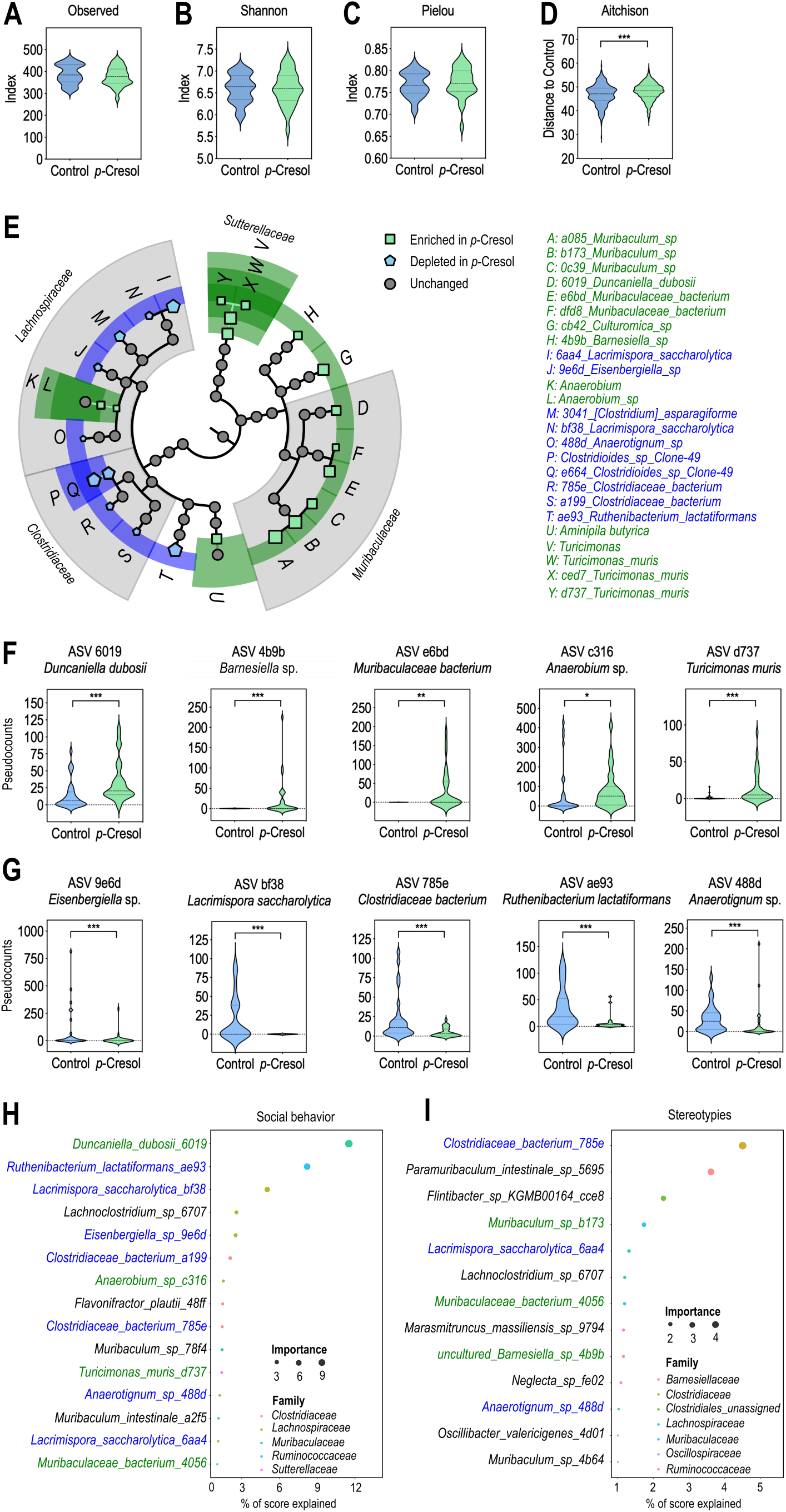
*p*-Cresol impacts microbial composition and selective bacterial taxa predict social deficits and stereotypies. (A-C) *α*-diversity as measured by Observed number of ASV (A), Shannon’s (B) and Pielou evenness’ (C) indexes; Kruskall-Wallis pairwise test: p>0.05 (n=30 Control, n=29 *p-*Cresol). (D) β-diversity as measured by Aitchison distances from Control and *p*-Cresol groups based on 16S rRNA gene sequencing. Group differences were tested by pairwise PERMANOVA, p=0.001 (n=30 Control, n=29 *p-*Cresol). (E) Synthetic cladogram presenting dysregulated ASV and bacterial taxa increased (green) or depleted (blue) upon *p*-Cresol exposure identified by ANCOM. Symbol size reflects the value of the corresponding CLR. (F, G) Pseudocounts plots from selected ASV increased (F) or depleted (G) upon *p-*Cresol exposure identified by ANCOM; Mann-Whitney U-Test: *p<0.05, **p<0.01, ***p<0.001 (n=30 Control, n=29 *p-*Cresol). (H, I) Microbial composition prediction of behavioral scores. ASV best predicting social behavior (H) or stereotypies (I) as identified by Random Forest analysis. Only ASV contributing >1% accuracy in behavioral scores prediction are presented. ASV identical to or ASV pointing towards the same species as ASV identified by ANCOM as increased or depleted upon *p*-Cresol exposure are labelled in green and blue respectively. (A-D, F, G) Data are presented as violin plots featuring frequency distribution of data, median (dashed line) and quartiles (dotted line).

### The microbiota from *p-*Cresol-treated mice induces social behavior deficits when transplanted to untreated recipients

To investigate whether microbiota composition changes may account for *p*-Cresol-induced behavioral alterations, we transplanted microbiota from either *p-*Cresol-treated or control mice to untreated recipients and assessed their behavior 3 weeks post-transplantation (Fig. 3A). For the sake of clarity, we will use FMT^Control^ as an abbreviation for “transplantation with the microbiota of control mice”. Likewise, we will refer to FMT*^p^*^-Cresol^ as an abbreviation for “transplantation with the microbiota of *p*-Cresol-treated mice”. Compared to FMT^Control^ mice, FMT*^p^*^-Cresol^ mice spent less time in social contact, exhibited decreased time, number, and mean duration of nose and paw contacts, but no change in the number of followings (Fig. 3B-G, Fig. S4A, B). Social deficits were accompanied by increased number of head shakes (Fig. 3H), but no changes in number of circling episodes (Fig. 3I), and of time spent grooming and digging (Fig. S4C, D). FMT^Control^ and FMT*^p^*^-Cresol^ groups were separated along the PC1 axis in PCA analysis of social interaction and stereotypies scores (Fig. 3J). Further, mice from these two groups behaved similarly in the novelty-suppressed feeding and zero-maze tests (Fig. S4E, F) suggesting that their anxiety levels were not impacted. To summarize, the transfer of microbiota from *p*-Cresol-treated mice to untreated recipients recapitulated the effects of *p*-Cresol treatment on social behavior deficits and partially on stereotypies.

**Fig. 3.**
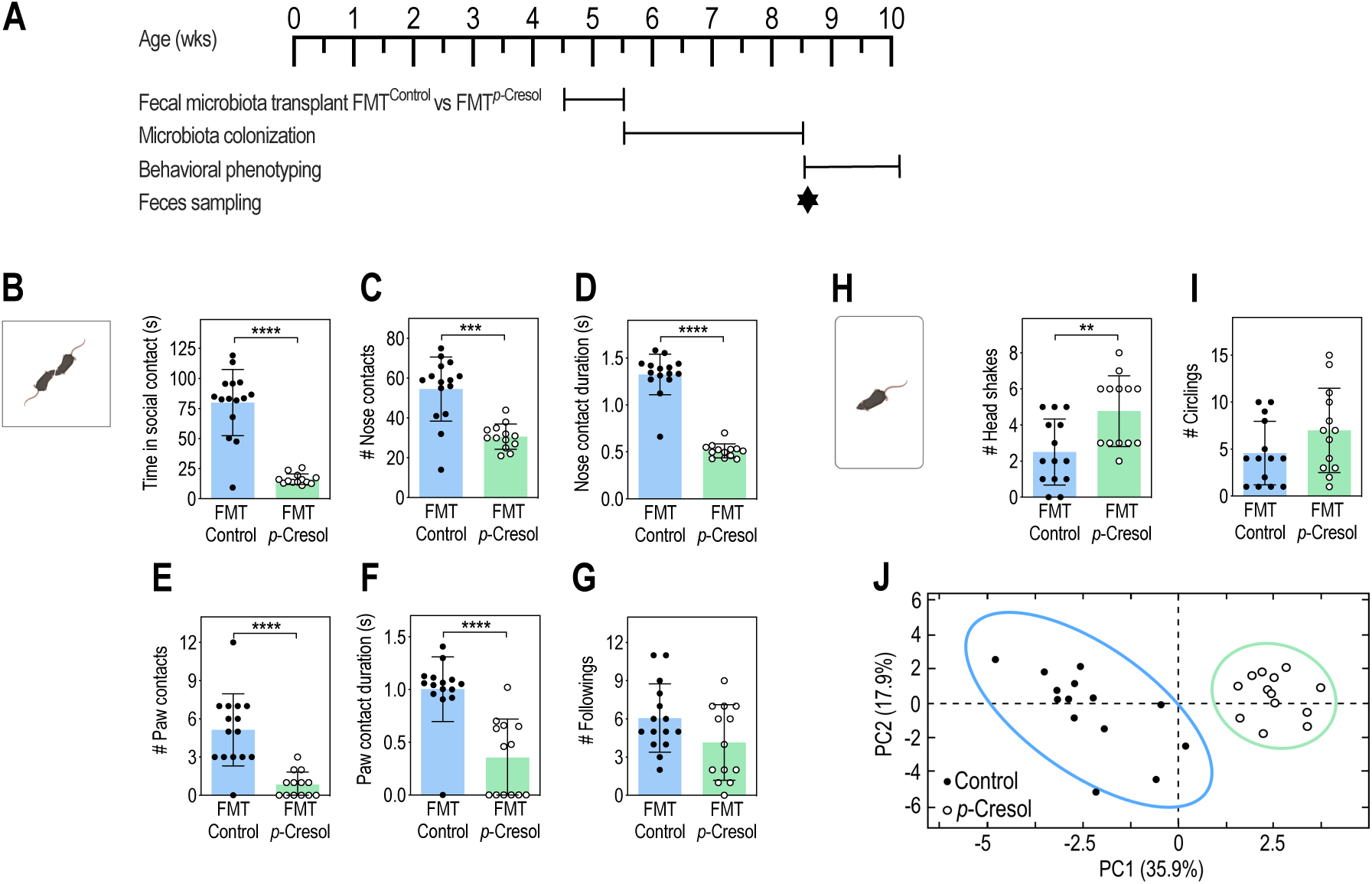
Microbiota remodeling following *p*-Cresol treatment induces social behavior deficits. (A) Timeline of the fecal microbiota transplant experiment from Control (FMT^Control^) or *p*-Cresol-treated donor animals (FMT*^p^*^-Cresol^) to recipient untreated mice. (B-G) Dyadic social interaction test (n=15 FMT^Control^, n=13 FMT*^p^*^-Cresol^): (B) Total time spent in social contact; Mann-Whitney U-test: ****p<0.0001. (C) Number of nose contacts; Mann-Whitney U-test: ***p<0.001. (D) Mean duration of nose contact; Mann-Whitney U-test: ****p<0.0001. (E) Number of paw contacts; Mann-Whitney U-test: ****p<0.0001. (F) Mean duration of each paw contact; Mann-Whitney U-test: ****p<0.0001. (G) Number of followings; Mann-Whitney U-test: p=0.1563. (H, I) Motor stereotypies: (n=14 FMT^Control^, n=13 FMT*^p^*^-Cresol^): (H) Number of head shakes; Mann-Whitney U-test: **p<0.01. (I) Number of circling events; Mann-Whitney U-test: p=0.1762. (J) PCA plots of behavioral scores recorded in the dyadic social interaction and direct monitoring of motor stereotypies (Figure 3B-I, Supplementary Figure 5I-L); ellipses of the 95% confidence intervals are indicated for each group (n=15 FMT^Control^, n=13 FMT*^p^*^-Cresol^). (B-I) Data are presented as dot-plots featuring means ± SD.

### Transfer of the microbiota from *p*-Cresol-treated mice to untreated recipients impacts the gut microbial ecology and increased fecal levels of *p*-Cresol

We used 16S rRNA sequencing to analyze the microbiota of FMT^Control^ and FMT*^p^*^-Cresol^ mice 3 weeks post FMT. Microbial richness and evenness were similar between the two groups as assessed by *α*-diversity indexes (Fig. 4A-C). However, analysis of β-diversity based on Aitchison’s distances revealed significant divergences in microbial composition between FMT^Control^ and FMT*^p^*^-Cresol^ mice (Fig. 4D), similarly to what was observed between *p*-Cresol-treated and control mice (Fig. 2D). ANCOM analysis revealed that 22 ASV, 5 species, 3 genera and 1 family were discriminant (|CLR|>0.2; W=0.7) (Fig. 4E, Tab. S2). Some of the dysregulated ASV related to taxa which we found as perturbed upon *p-*Cresol exposure (presented in Fig. 2E, F, Tab. S1). Notably, ASV counts related to several *Muribaculaceae* (*Duncaniella* sp. B8, *Duncaniella dubosii*, one *Barnesiella* sp.) and one *Anaerobium* sp. were increased in FMT*^p^*^-Cresol^ mice (Fig. 4E, F, Tab. S2). In addition, ASV counts related to one *Eisenbergiella* sp., *Lacrimispora saccharolytica,* and one *Clostridiaceae* bacterium were decreased in FMT*^p^*^-Cresol^ mice (Fig. 4E, G, Tab. S2). Further, these ASV contributed to social abilities prediction in FMT^Control^ and FMT*^p^*^-Cresol^ mice, with notably one *Barnesiella* sp., *Duncaniella dubosii,* and one *Anaerobium* sp. explaining 8%, 6% and 4% of the score, respectively (Fig. 4H). Also, several of these ASV overlapped with the ASV already identified as predictors of social behavior in control and *p*-Cresol-treated mice (Fig. 2H). This latter result suggested that specific bacterial taxa could be responsible for social deficits induced by both *p*-Cresol treatment and FMT*^p^*^-Cresol^. This was not the case for stereotypies (Fig. 4I), which were only modestly explained by microbiota composition post FMT^Control^ and FMT*^p^*^-Cresol^.

**Fig. 4.**
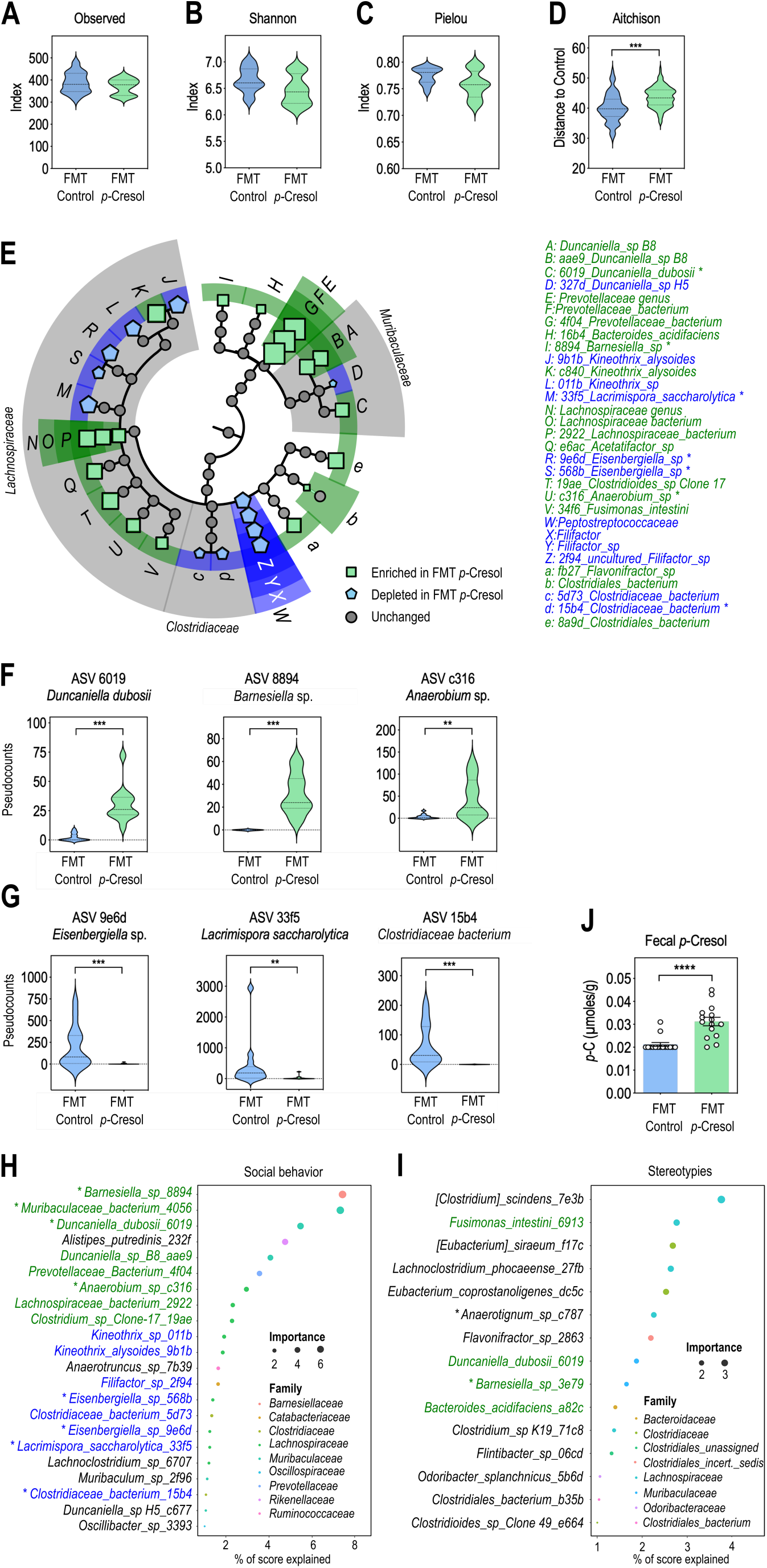
Transfer of *p*-Cresol-induced microbiota impacts microbial composition, increases *p*-Cresol production and selective bacterial taxa predict social deficits and stereotypies. (A-C) *α*-diversity as measured by Observed number of ASV (A), Shannon’s (B) and Pielou evenness’ (C) indexes; Kruskall-Wallis pairwise test: p>0.05 (n=14 FMT^Control^, n=12 FMT*^p-^*^Cresol^). (D) β-diversity as measured by Aitchison distances from Control and *p*-Cresol groups based on 16S rRNA gene sequencing. Group differences were tested by pairwise PERMANOVA, p=0.001 (n=14 FMT^Control^, n=12 FMT*^p-^*^Cresol^). (E) Synthetic cladogram presenting dysregulated ASV and bacterial taxa associated with FMT*^p^*^-Cresol^ (green) or FMT^Control^ class (blue) identified by ANCOM. Symbol size reflects the value of the corresponding CLR. Asterisks indicate ASV identical to (or ASV with similar affiliations) to ASV also dysregulated upon *p*-Cresol exposure (Fig. 2E). (F, G) Pseudocounts plots from selected ASV increased (F) or depleted (G) upon FMT*^p-^*^Cresol^ identified by ANCOM; Mann-Whitney U-Test: **p<0.01, ***p<0.001. (H, I) Microbial composition prediction of behavioral scores after FMT (n=14 FMT^Control^, n=12 FMT*^p-^*^Cresol^). ASV best predicting social interaction deficits (H) and stereotypies (I), as identified by Random Forest analysis. Only ASV contributing >1% accuracy in behavioral scores prediction are presented. ASV identical to or ASV pointing towards the same species as ASV identified by ANCOM as increased or depleted in FMT*^p-^*^Cresol^ as compared to FMT^Control^ are labelled in green and blue respectively. Asterisks indicate ASV identical to (or ASV with similar affiliations) to ASV also predicting behavioral scores upon *p*-Cresol exposure (Fig. 2H, I). (J) Fecal levels of *p*-Cresol 3 weeks post FMT (n=15 FMT^Control^, n=15 FMT*^p-^*^Cresol^); Mann-Whitney U-test: ****, p<0.0001. (A-D, F, G) Data are presented as violin plots featuring frequency distribution of data, median (dashed line) and quartiles (dotted line). (J) Data are presented as dot-plots featuring means ± SD.

In addition, FMT*^p^*^-Cresol^ mice exhibited increased fecal levels of *p*-Cresol (Fig. 4J), suggesting that the microbiota of these mice produced more *p-*Cresol compared to the one from FMT^Control^ mice. We therefore searched for proteins homologous to enzymes involved in *p*-Cresol biosynthesis in bacterial taxa presenting increased abundance in both *p*-Cresol-treated mice and FMT*^p^*^-Cresol^ (*Duncaniella dubosii, Barnesiella* spp. and *Anaerobium* spp.). Two microbial metabolic pathways are involved in the production of *p-*Cresol from tyrosine. The direct pathway involves the tyrosine lyase ThiH described in *Escherichia coli* [38] (Fig. S5A). The indirect pathway involves several enzymes, with the tyrosine aminotransferase TyrB catalyzing the initial step and the *p*-hydroxyphenylacetate decarboxylase Hpd A, B and C subunits ([28] described in *Clostridioides difficile*) catalyzing the final step (Fig. S5A). Our Blast analysis revealed that ThiH homologous proteins were present in *Anaerobium acetethylicum*, and several *Muribaculaceae* members: *Duncaniella dubosii* and *Barnesiella* spp. and in (Tab. 1, Tab. S3). Regarding the indirect pathway, we only identified enzymes homologous to HpdA in *Anaerobium acetethylicum* and *Duncaniella dubosii,* but with generally lower identity scores, and no homolog for TyrB or HpdB/C (Tab. 1, Tab. S3). This suggested that the direct pathway was likely privileged for *p-*Cresol synthesis in those bacterial species.

To confirm bacterial species possibly accounting for *p*-Cresol production post FMT^Control^ and FMT*^p^*^-Cresol^, we used RF analysis to identify the ASV counts best predicting fecal *p-*Cresol levels. Several ASV related to *Clostridioides* spp. and one *Anaerostipes* sp. were identified (Fig. S5), in line with previous studies which identified relatives of these taxa as bacterial *p*-Cresol producers [28]. In addition, four *Muribaculaceae* spp., incl. *Duncaniella dubosii* and one *Barnesiella* sp. contributed to accurately predict *p*-Cresol fecal levels (Fig. S5), reinforcing the possible link between *p*-Cresol production and these species overabundant in both *p*-Cresol-treated and FMT*^p^*^-Cresol^ mice.

### Transfer of a normal microbiota to *p*-Cresol-treated recipients restores social behavior, VTA dopamine neurons excitability and fecal *p*-Cresol

Having shown that FMT*^p^*^-Cresol^ mice exhibited social behavior deficits, we investigated whether *p-*Cresol-induced behavioral alterations were restored by FMT^Control^ (Fig. 5A). While *p-*Cresol-treated mice displayed both social behavior deficits (Fig. 5B-H) and stereotypies (Fig. 5J,K), as already shown (Fig. 1B-N, Fig. S3B-O), FMT^Control^ to *p*-Cresol-treated mice normalized both the number, duration (Fig. 5B, C, E, G, Fig. S6I, J) and quality of social contacts (Fig. 5D, F) in the dyadic social interaction test. The sociability index (Fig. 5H), the preference for the interactor mouse (Fig. S6G, H), and the quality of social interactions (Fig. 5I) were also rescued as assessed with the three-chamber test. While social deficits were fully restored, stereotyped behaviors were only partially normalized (Fig. 5J, K, Fig. S6K, L). Normalization of *p-*Cresol-induced behaviors by FMT^Control^ was confirmed by PCA analysis with control mice pre- and post FMT^Control^ and *p-*Cresol-treated mice post FMT^Control^ clustered on the right side of the PC1 axis, while *p*-Cresol-treated mice stood alone on the left side (Fig. 5L). Most importantly, FMT^Control^ to *p*-Cresol-treated mice restored both the excitability of VTA dopamine neurons (Fig. 5M, N) and fecal *p*-Cresol levels (Fig. 5O).

**Fig. 5.**
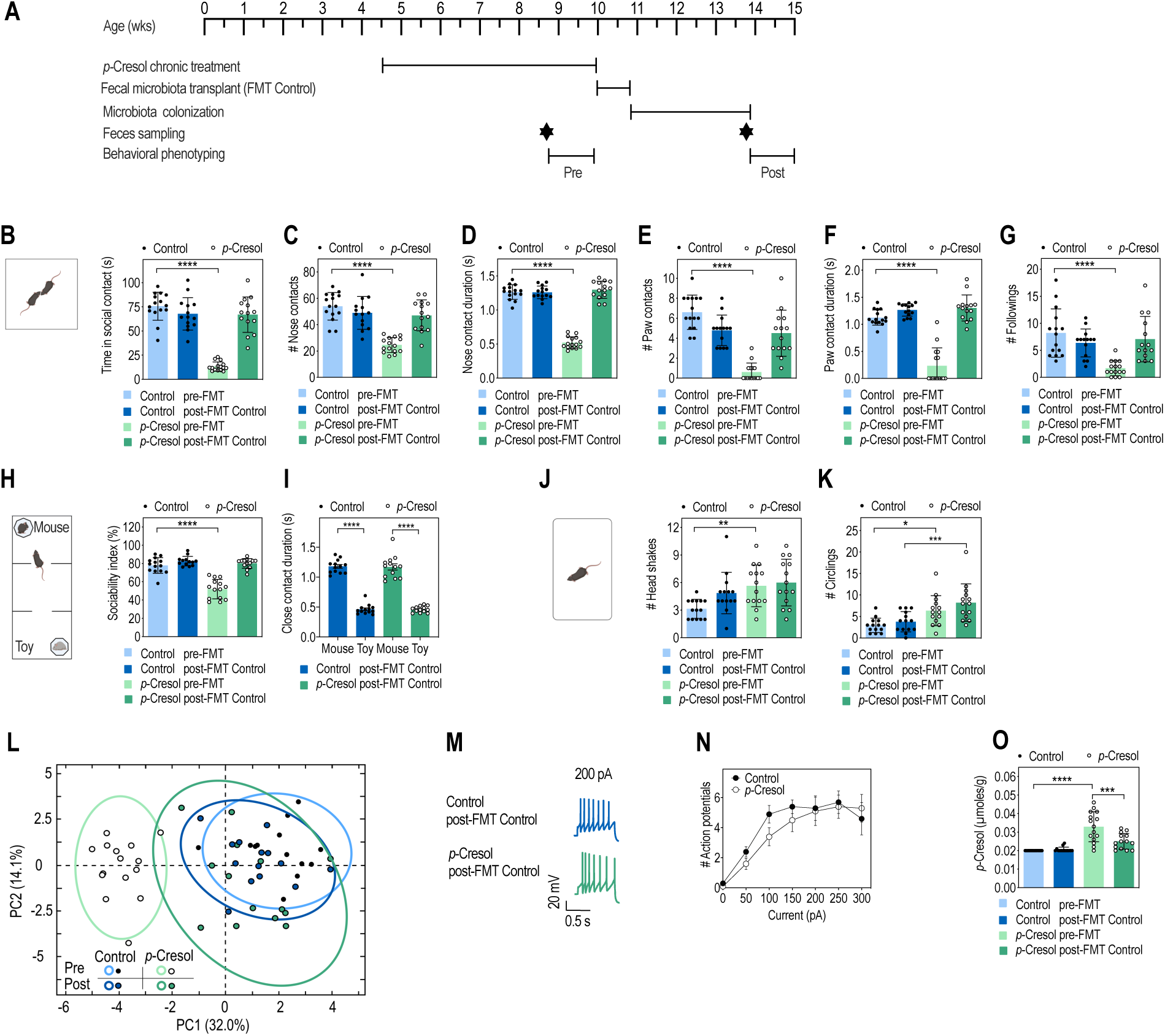
Transfer of microbiota from control mice to *p*-Cresol-treated mice restores social behavior deficits, VTA dopamine neurons excitability and fecal *p*-Cresol levels. (A) Timeline of the fecal microbiota transplant experiment of fecal slurry from control donor mice (FMT^Control^) to control or *p*-Cresol-treated recipient mice. (B-G) Dyadic social interaction test pre-FMT^Control^ and 3 weeks post-FMT^Control^ (n=15/group pre-FMT, n=14/group post-FMT): (B) Total time spent in social contact; 2-way ANOVA: p(Treatment)<0.0001, p(FMT^Control^)<0.0001, p(Treatment x FMT^Control^)<0.0001; Šidák’s post-hoc tests for treatment effect: ****p<0.0001 for pre-FMT groups, p>0.05 for post- FMT groups. (C) Number of nose contacts; 2-way ANOVA: p(Treatment)<0.0001, p(FMT^Control^)= 0.0022, p(Treatment x FMT^Control^)<0.0001; Šidák’s post-hoc tests for treatment effect: ****p<0.0001 for pre-FMT groups, p>0.05 for post- FMT groups. (D) Mean duration of each nose contact; 2-way ANOVA: p(Treatment)<0.0001, p(FMT^Control^)<0.0001, p(Treatment x FMT^Control^)<0.0001; Šidák’s post-hoc tests for treatment effect: ****p<0.0001 for pre-FMT groups, p>0.05 for post- FMT groups. (E) Number of paw contacts; 2-way ANOVA: p(Treatment)<0.0001, p(FMT^Control^)=0.0221, p(Treatment x FMT^Control^)<0.0001; Šidák’s post-hoc tests for treatment effect: ****p<0.0001 for pre-FMT groups, p>0.05 for post- FMT groups. (F) Mean duration of each paw contact; 2-way ANOVA: p(Treatment)<0.0001, p(FMT^Control^)<0.0001, p(Treatment x FMT^Control^)<0.0001; Šidák’s post-hoc tests for treatment effect: ****p<0.0001 for pre-FMT groups, p>0.05 for post- FMT groups. (G) Number of followings; 2-way ANOVA: p(Treatment)=0.0020, p(FMT^Control^)=0.0516, p(Treatment x FMT^Control^)=0.0002; Šidák’s post-hoc tests for treatment effect: ****p<0.0001 for pre-FMT groups, p>0.05 for post- FMT groups. (H, I) 3-chamber test pre-FMT^Control^ and 3 weeks post-FMT^Control^ (n=15/group pre-FMT, n=14/group post-FMT): (H) Sociability index; 2-way ANOVA: p(Treatment)=0.0004, p(FMT^Control^)<0.0001, p(Treatment x FMT^Control^)=0.0032; Šidák’s post-hoc tests for treatment effect: ****p<0.0001 for pre-FMT groups, p>0.05 for post-FMT groups. (I) Mean duration of each nose contact with the mouse interactor or toy mouse; 2-way ANOVA: p(FMT^Control^)=0.9067, p(Mouse-Toy)<0.0001, p(FMT^Control^ x Mouse-Toy)=0.8713; Šidák’s post-hoc tests for Mouse vs. Toy preference: ****p<0.0001. (J, K) Motor stereotypies pre-FMT^Control^ and 3 weeks post-FMT^Control^ (n=15/group pre-FMT, n=15/group post-FMT): (J) Number of head shakes; 2-way ANOVA: p(Treatment)=0.0021, p(FMT^Control^)=0.0715, p(Treatment x FMT^Control^)=0.2334; Šidák’s post-hoc tests for treatment effect: **p<0.01 for pre-FMT groups, p>0.05 for post-FMT groups. (K) Number of circling events; 2-way ANOVA: p(Treatment)<0.0001, p(FMT^Control^)=0.1133, p(Treatment x FMT^Control^)=0.5555; Šidák’s post-hoc tests: *p<0.05 for pre-FMT groups, ***p<0.001 for post-FMT groups. (L) PCA plots of behavioral scores recorded in the dyadic social interaction and direct monitoring of motor stereotypies pre-FMT^Control^ and 3 weeks post-FMT^Control^ (Figure 5B-G, J, K, Supplementary Figure 6I-L); ellipses of the 95% confidence intervals are indicated for each group (n=15/group pre-FMT, n=14/group post-FMT). (M, N) Electrophysiological recordings of dopamine neurons activity in the VTA 3 weeks post-FMT^Control^: (M) Representative traces of dopamine neurons activity in patch-clamp experiments performed post-FMT^Control^ after a 200 pA current injection. (N) Number of action potentials evoked by different current injection steps (n=3 animals/group, 10 cells recorded/animal); 2-way ANOVA: p(FMT^Control^)=0.5474, p(Current)<0.0001, p(FMT^Control^ x Current)=0.3640; Šidák’s post hoc tests for treatment effect: p>0.05. (O) Fecal levels of *p*-Cresol pre- and 3 weeks post-FMT^Control^ (n=15/group pre-FMT, n=14/group post-FMT); 2-way ANOVA: p(Treatment)<0.0001, p(FMT^Control^)=0.0044, p(Treatment x FMT^Control^)=0.0011; Šidák’s post hoc tests: ****p<0.0001; ***p<0.001. (B-K, O) Data are presented as dot-plots featuring means ± SD. (N) Data are presented as means ± SD.

## DISCUSSION

### p-Cresol selectively induces ASD core behavioral symptoms

Here we show that *p*-Cresol-treated mice exhibit social behavior deficits and stereotypies, but no changes in anxiety, locomotion, or cognition. This suggests a possible causal relationship between elevated *p*-Cresol levels and ASD core symptoms. While several other metabolites modified behavior when administered to rodents, none of them selectively induced ASD core symptoms. For example, the SCFA propionate did not only induce social interaction deficits and stereotypies, but also anxiety and hyperlocomotion [16]. Indoles increased social contacts and anxiety, reduced locomotor activity in rats [39] and exacerbated emotional behaviors in chronically stressed mice [40]. 4-EPS increased anxiety and startle reflex in mice, but had no impact on social behavior or stereotypies [19]. Frequent ASD comorbidities are ADHD, ID and anxiety disorder [3] and we show that related behavioral domains (locomotor activity, cognition, anxiety) are not impacted by *p*-Cresol exposure in mice. It appears that propionate, indoles or 4-EPS induce anxiety, which could interfere with social and cognitive abilities and explain their broad effects. These data collectively suggest that microbial metabolites likely interfere with several dimensions of behavior impacted in ASD, each with its specificities, with *p*-Cresol selectively impacting social behavior and stereotyped/perseverative behaviors.

Lastly, a recent study showed that acute intravenous (i.v.) *p*-Cresol injection exacerbated anxiety, hyperactivity, stereotypies and reduced social preference in the BTBR model of idiopathic ASD [41]. Compared to the selective effects of *p*-Cresol in our model, these broader effects may be explained by differences in genetic background (BTBR versus C57BL/6) or the mode of *p-*Cresol administration (i.v. versus oral, acute versus chronic). Since primary exposure to microbial metabolites occurs in the GI tract, the *per os* delivery of *p*-Cresol that we used reflects more closely than i.v. GI exposure to endogenous *p*-Cresol produced by the microbiota. We have shown that this does not lead to elevated circulating *p-*Cresol, suggesting limited systemic effects compared to i.v. injection, and possibly explaining the selective induction of social deficits and stereotyped/perseverative behaviors.

### p-Cresol induces behavioral impairments via microbiota remodeling

Three independent meta-analyses reported bacterial β-diversity changes associated with increased abundance of the genus *Clostridioides* and decreased abundance of the genus *Bifidobacterium* in ASD patients as compared to neurotypical controls [7–9]. While we observed β-diversity changes in *p-*Cresol-treated mice, *Clostridioides* and *Bifidobacterium* were not impacted at the genus level. Instead, we observed selective changes in lower taxonomic levels (mostly ASV, species). Confounding factors (diet, treatment, sex, age) have not been considered in most human microbiota studies, possibly leading to discrepancies in the identified changes [7–9]. Also, the human and murine microbiota differ in terms of bacterial species representation. Indeed, we highlighted selective changes in murine-specific bacteria upon *p*-Cresol exposure. Notably *Muribaculaceae* and *Turicimonas* are both dominant taxa in the murine microbiota [42]. While changes in *β−*diversity have been observed in ASD murine models [19–23], the lower taxonomic resolution of most studies precludes adequate comparison with our findings in *p*-Cresol-treated mice. However, it is worth mentioning that mouse and human microbiota share 95.2% of metabolic functions [43], we therefore posit that studying *p*-Cresol, which is a product from microbial metabolism common to murine and human microbiota, reinforces the translationality of our findings. Our data therefore suggest that *p*-Cresol could contribute to ASD core symptoms in humans.

Two metabolic pathways have been described for *p-*Cresol production from tyrosine. The direct pathway described in *E. coli* involves ThiH [38], while the indirect pathway present in *C. difficile* and *Blautia hydrogenotrophica* involves both TyrA/B/C and Hpd [28]. These enzymes have not yet been formally described in *Muribaculaceae*, such as *Duncaniella dubosii* or *Barnesiella* spp., or in *Anaerobium* spp. (which are upregulated taxa in *p-*Cresol-treated and FMT*^p^*^-Cresol^ mice), but ThiH homologous proteins are detected, suggesting that they could produce *p-* Cresol via the direct pathway. The possible ability of *Muribaculaceae* members to synthetize *p*-Cresol is reinforced by the fact that the counts of ASV related to *Duncaniella dubosii*, one *Barnesiella* sp. and several *Muribaculaceae* bacteria predict fecal *p-*Cresol levels in FMT*^p^*^-Cresol^ mice. *Muribaculaceae* have been recently described and functional analysis have revealed their ability to metabolize benzoate, a known degradation product of *p-*Cresol [44]. Future work is required to understand their metabolic capacities.

Conversely, FMT^Control^ to *p-*Cresol-treated mice decreased *p-*Cresol levels. This latter result suggested that the capacity to metabolize *p-*Cresol was reduced after FMT^Control^, possibly explaining the rescue of both social behavior and VTA dopamine neurons excitability. Finally, the fact that FMT*^p^*^-Cresol^ and FMT^Control^ respectively induce or rescue social deficits, and to a lesser extent stereotypies, argues for a strong contribution of the *p*-Cresol-induced changes in microbial composition to social behavior in our model. Future metagenomic studies will help refine the bacterial species and metabolic pathways linking *p*-Cresol to social behavior deficits.

### Exposure to p-Cresol impacts central dopamine neurons activity

Social interactions are pleasurable events for humans and animals, as shown by the activation of the reward circuit by social stimuli. This activation is blunted in ASD [45]. The VTA is a key subcortical node of the mesolimbic reward pathway [46] and ASD patients display defects in VTA connectivity [31]. Moreover, blocking VTA dopamine neurons reduced social interactions in rodents [30, 47]. Further, impaired VTA dopamine neurons activity was observed in both environmental (MIA, DIO) [20, 35] and genetic (*Shank3b-*, *Nlgn3-* and *Ube3a-*deficient mice) [21, 32–34] mouse models of ASD. The reward system and associated dopamine circuits are influenced by the microbiota [48]. Gut-to-brain neural circuit establishes vagal neurons as essential components of the central reward pathway, linking sensory neurons in the gut to central dopamine release by the substantia nigra, a brain area known to project to the VTA [49]. Microbiota manipulations normalize both social behavior and VTA DA neurons activity in *Shank3b-*deficient and DIO ASD models [20, 21], as well as in *p*-Cresol-treated mice (this study). Taken together, these results support a model in which *p*-Cresol-induced changes in microbial composition impair VTA dopamine neurons activity, possibly disrupting dopamine in the reward circuit, and contributing to social behavior deficits. Our data are in agreement with a recent study showing that acute intravenous administration of *p*-Cresol increased dopamine turnover in several areas of the reward circuit in the BTBR ASD model [41]. How *p-*Cresol-induced microbiota changes impacts VTA activity, and possibly central dopamine and the social reward circuit remains to be further investigated.

Some microbial metabolites are ligands to host receptors: 3-indoxylsulfate binds to the aryl-hydrocarbon receptor [50], indole-3-propionate to the xenobiotic sensor pregnane X receptor [51] and propionate to GPR41 and GPR43 [52]. Yet, their signaling role has been mainly investigated in metabolic, GI or autoimmune disorders [50–52], and not in ASD. Indoles, SCFA and their receptors are detected in the brain suggesting that these metabolites could act centrally [25, 53, 54], a hypothesis that remains to be tested. While both urinary and fecal *p*-Cresol levels are increased in *p-*Cresol-treated mice, serum levels are not, suggesting that *p-* Cresol does not reach bioactive levels in brain. In contrast, the results of our FMT experiments demonstrated that *p-*Cresol triggers ASD-like behaviors by remodeling the microbiota. The microbiota induced by *p*-Cresol could influence the activity of gut immune and neuroendocrine cells, as well as of vagal afferences, resulting in perturbations of the microbiota gut-brain axis and altered behavior [1, 25]. How the effects of *p*-Cresol-induced microbiota remodeling are relayed to the brain to modulate social behavior remains to be investigated.

### Conclusions and perspectives

We found that *p*-Cresol-treated mice exhibited social interaction deficits and stereotypies, reminiscent of ASD core symptoms in humans. These behaviors were dependent on changes in microbiota composition and were associated with reduced excitability of VTA dopamine neurons as well as elevated *p-*Cresol production. Because *p*-Cresol levels are elevated in ASD patients, our study suggests that increased levels of this metabolite could contribute to ASD core symptoms in humans. Further, the ability of a control microbiota to normalize *p-*Cresol levels, VTA dopamine neurons excitability and social behavior when transplanted to *p-*Cresol-treated mice provides a rationale for clinical trials aimed at studying the beneficial impact of microbiota interventions targeting *p*-Cresol production to alleviate core ASD symptoms and in particular social deficits.

## METHODS

### Ethical considerations

Animal housing and experimentation were conducted in facilities certified by regional authorities (Direction Départementale de Protection des Populations des Alpes-Maritimes, accreditation #EC-06-152-5). The procedures involved in this study were approved by the local ethics committee for animal experimentation (Ciepal-Azur) and the Ministère de l’Enseignement Supérieur et de la Recherche, in agreement with the European Communities Council Directive (2010/63EU) for animal experiments.

### Animals treatment

Weaned C57BL/6J male mice (21 to 28 day-old) were ordered to Charles River (France). Animals were randomly assigned to experimental groups and housed in medium-size (up to 5 animals) open cages filled with wooden bedding, one plastic house and nesting material for enrichment, in a temperature-(22-24°C) and hygrometry-(70-80%) controlled room, under 12 hours light–dark cycle (8:00 a.m.-8:00 p.m.). Mice had *ad libitum* access to water and standard chow (reference 4RF25, Mucedola). Mice were habituated for 6 days to our animal facility before starting treatment with *p*-Cresol (reference W233706-SAMPLE-K, Sigma-Aldrich) dispensed in sterile drinking water at a concentration of 0.25 g/L. Bottles were renewed twice weekly. This amounts to a dose of 50 mg/Kg/24 h based on a mean body mass of 25 g and a mean drinking water consumption of 5 mL/24 h. Body weight, consumption of water and chow were monitored weekly.

### Quantification of *p-*Cresol in urine and serum by gas chromatography coupled to mass spectrometry (GC-MS)

Mice were individually placed in a clean empty plastic cage and urine was collected over 10 min and centrifuged (12 000 g, 5 min). The supernatant was snapped-frozen in liquid nitrogen and stored at −80°C until analysis. Blood was collected by cardiac puncture, transferred in Eppendorf tubes and allowed to clot on bench RT for 30 min. After centrifugation (12000g, 10 min, RT), serum was collected, snapped-frozen in liquid nitrogen and stored at −80°C until analysis. Prior to gas chromatography-mass spectrometry (GC-MS), urine (20 μL) and serum (100 μL) samples were prepared as follows: i) samples were spiked with 10 μL internal standard solution (myristic acid-d_27_ in isopropanol, 750 mg/mL), ii) 850 μL of ice cold methanol were added, followed by centrifugation for 20 min (4°C, 16000 g), iii) 750 μL of supernatants were transferred to silanized dark 2 mL autosampler vials and evaporated to dryness in a rotational vacuum concentrator (45°C, 20 mbar, 2 h), iv) 50 μL of methoxyamine solution (2% in pyridine) were added and the samples were incubated overnight at RT, and v) 100 μL of N-methyl-trimethylsilyl-trifluoroacetamide (MSTFA) containing 1% of trimethylchlorosilane (TMCS) solution were added, the samples were incubated at 60°C for 1 h and transferred to dark autosampler vials with 250 μL silanized inserts. The detailed instrumental conditions for GC-MS analysis were based on a previous study [55], with minor modifications to quantify *p-*Cresol. Briefly, the samples were analyzed in an Agilent 7890B-5977B Inert Plus GC-MS system. Two μL of each sample were injected in a split inlet (1:10 ratio). The chromatographic column was an Agilent ZORBAX DB5-MS (30 m X 250 µm X 0.25 µm + 10 m, Duraguard). The temperature gradient was 37.5 min long and the mass analyzer was operated in full scan mode between 50-600 m/z. Blank and pooled quality control (QC) samples in several dilutions were included in the analysis to ensure that the measurements were reproducible. There was no external contaminations and the monitored analytes were responding linearly to the detector.

### Quantification of *p*-Cresol in fecal samples by liquid chromatography-tandem mass spectrometry (LC-MS/MS)

Mice were individually placed in a clean empty plastic cage and feces were collected over 10 min and immediately snapped-frozen in liquid nitrogen and stored at −80°C until analysis. Thawed fecal pellets were extracted as previously described in [56]. Briefly fecal pellets were homogenized in ice-cold water (1:1 w:v ratio), vortexed 30 s, ultrasonicated for 2 min, then subjected to ultracentrifugation (175,000 g, 30 min, 4 °C). For the determination of unconjugated *p-*Cresol by LC-MS/MS, fecal supernatant preparation included addition of an internal standard (1 ng/mL of *p*-Cresol-d8, Eurisotop, St-Aubin, France) and derivatization with dansyl chloride according to [57]. Dansylation was shown to improve signal intensity in LC-MS/MS analysis of low-abundance phenolic compounds [58]. Briefly, 100 µL of fecal supernatant were mixed with 300 µL of an acetonitrile solution vortexed, incubated at −20°C for 20 min and centrifuged (12,000 g, 7 min, 4°C). Two hundred µL of the resulting supernatant were mixed with 50 µL of 0.1 M carbonate-bicarbonate solution (pH = 10), 125 µL of water, and 125 µL of dansyl-chloride at a final concentration of 0.5 mg/mL. After vortexing, the mixture was incubated for 10 min at 60°C and extracted with 2.5 mL of hexane. Hexane residues were air-dried and reconstituted with 1 mL of acetonitrile-water (1:1, v/v). Fifteen µL of sample were injected and analyzed using a Waters ACQUITY ultraperformance liquid chromatography (UPLC) system equipped with a binary solvent delivery manager and sample manager (Waters Corporation, Milford, MA, USA) and coupled to a tandem quadrupole-time-of-flight (Q-TOF) mass spectrometer equipped with an electrospray interface (Waters Corporation). Quantifications were performed by referencing calibration curves obtained with internal standards. Unconjugated *p*-Cresol was identified by comparing with the accurate mass and the retention time of the reference standard (*p*-Cresol, Sigma-Aldrich) in our in-house library.

### Behavioral testing

Behavioral phenotyping started after a 4-week treatment with *p*-Cresol and lasted for 1-2 weeks during which treatment was continued. All behavioral experiments were performed during the day. The animals followed multiple behavioral tests respecting their resting hours, with a minimum of 2 days between tests. Standard behavioral tests that have extensively been used to characterize genetic and environmental models of ASD [59–63] were implemented to assess the behavioral dimensions impacted in ASD.

### Social abilities

#### Three-chamber sociability test

The stimulus mouse was habituated to the cylindrical cages (20 min sessions, repeated 3 days before the testing day). The apparatus consisted in a rectangular non-transparent plexiglass box (60×30 cm), divided in three compartments (object chamber, empty chamber and unfamiliar mouse chamber) connected by open doors (4 cm width) to allow the mouse to move freely between the different compartments. The unfamiliar mouse or the toy mouse were placed in wired cylindrical cages. The test consisted in a habituation phase to the empty apparatus only in the presence of empty cylindrical cages during 5 min and a sociability phase where the unfamiliar mouse or the toy mouse is introduced in the cylindrical cages. Time spent in the 2 chambers were video-tracked over 10 min and analyzed *a posteriori* during the habituation and sociability phase using the ANY-maze software. The number and the time spent in close contact with each of the empty cylindrical cage (during the habituation phase) as well as with the stimulus mouse or with the toy mouse contained in wired cylindrical cage (during the sociability phase) were manually scored by an experienced experimenter blind to the experimental group. The mean duration of close contacts was calculated from these data [60, 64–66]. The relative position of stimulus unknown mouse (versus mouse toy) was counterbalanced between groups. There was no bias in chamber preference, empty cylindrical cages preference or interaction ratio during the habituation phase (Fig. S2A-E, Fig. S6A-E).

#### Dyadic social interactions

Direct social interactions were recorded in open-field arena with a low light intensity (15 Lux). The subject mouse was put in presence with un unfamiliar sex- and age-matched interactor and their interaction was recorded for 10 min. Manual scoring by an experienced experimenter blind to the experimental group was performed *a posteriori* by recording time spent in social contact events, number of nose and paw contacts, time spent in nose and paw contact, number and time spent in self-grooming, number of self-grooming events after social contact, number of rearing and grooming events, number of circling episodes and head shakes. The mean duration of nose and paw contacts was calculated from these data [60, 64–66].

### Stereotypic/perseverative behaviors

#### Motor stereotypies

The subject mouse was placed in a clean standard home cage (21×11×17 cm) covered with a thick layer of fresh sawdust (4 cm) and recorded for 10 min with light intensity set at 40 Lux. Manual scoring by an experienced experimenter blind to the experimental group was performed *a posteriori* by computing the number of events of head shake, rearing, digging, grooming, circling episodes and total time spent digging and grooming [59].

#### Marble burying test

Marble burying was used as a measure of perseverative behavior [67]. The subject mouse was placed in a clean standard home cage (21×11×17 cm) filled with 4 cm of fresh sawdust on which 20 glass marbles (diameter 1.5 cm) were disposed with light intensity set at 40 Lux. The number of marbles buried (more than half of its surface outside of sawdust) during a 30 min session was monitored by an experienced experimenter blind to the experimental group [59].

#### Y maze spontaneous alternation task

Spontaneous alternation behavior was used to assess behavioral flexibility and perseveration [68–70]. The apparatus consisted in three arms made of plexiglass (40×9×16 cm) containing different symbols to be differentiated. We placed the mice in the centre and allowed it to explore the arms for 5 min with a low light intensity of 15 Lux. We measured the willingness of mice to explore new environments and we scored different patterns as spontaneous alternation (SPA), alternate arm returns (AAR) and same arm return (SAR). We then calculated the percentage of SPA, AAR and SAR[59, 71].

### Locomotor activity

#### Actimetry

The measurement of individual spontaneous activity was performed in actimetry chambers (Imetronic) consisting of cages equipped with infrared beams able to detect in real time horizontal and vertical movements (rearing events). Animals were individually placed in actimetry chambers under a 12-hr light/dark cycle, with free access to food and drinking water. To avoid biases in measurements due to stress possibly inducing hyperlocomotion in a novel environment, a habituation period (Day 0: 11:00 p.m. - Day 1: 8:00 a.m.) preceded the 24 h-recording of horizontal activity (Day 1: 8:00 a.m. – Day 2: 8:00 a.m.)[61].

### Anxiety

#### Open field test

Mice were individually placed in the corner of a white and opaque quadratic arena (40×40 cm) and allowed to explore for 10 min under low illumination (15 Lux). Total distance travelled, time spent in center and number center entries were analyzed by videotracking using the ANY-Maze software [61, 62].

#### Novelty suppressed feeding test

Mice were food-deprived for 16 h, with unlimited access to drinking water. A pellet of food was positioned in the center of an open-field covered with 1 cm of sawdust with light intensity set at 60 Lux. The subject mouse was then introduced in a corner of the arena and the latency (s) for the first bite in the pellet recorded. Immediately after the test, the animal was placed in a clean cage with *ad libitum* access to food and water [59, 71]. *Zero-maze.* The zero-maze apparatus (56 cm diam.) is divided in four equal quadrants with two opposing open quadrants and two opposing closed quadrants with grey acrylic walls (13 cm height). At the beginning of the trial, the subject mouse was placed head facing the entrance of a closed arm, and allowed to explore the ring for 5 min under high light intensity (200 Lux). The time spent in the open arms was scored *a posteriori* on videorecordings by a trained experimenter.

### Cognition

#### Novel object recognition test

The novel object recognition test was performed in a rectangular arena (20×40 cm). The test consisted of three sessions under constant light intensity (15 Lux). During the habituation session, the mice explored freely the empty arena for 5 min. For the training session, two identical objects were placed on each side of the arena and the mouse allowed to explore for 10 min. For the test session, one of the objects was replaced by a novel object and the mouse allowed to explore the objects for 10 min. The time spent exploring the familiar and novel objects by the subject animal were manually scored *a posteriori* on videorecordings of the sessions. A recognition index was calculated as the percentage of the time spent exploring the novel object over the total time spent exploring both objects [72].

### *Ex vivo* patch-clamp electrophysiological recordings

Electrophysiological recordings were performed on sub-groups of control or *p*-Cresol-treated which had not been subjected to behavioral tests but nevertheless belonged to larger cohorts in which behavioral impairments in the *p-*Cresol group were observed. Mice were anesthetized (Ketamine (150 mg/kg)/ Xylazine (10 mg/kg), trans-cardially perfused with artificial cerebrospinal fluid (aCSF solution: NaCl 119 mM, KCl 2.5 mM, NaH_2_PO_4_ 1.25 mM, MgSO_4_ 1.3 mM, CaCl_2_ 2.5 mM, NaHCO_3_ 26 mM, glucose 11 mM). Brains were removed from skull and sliced sagittally in 250 μm sections using a HM650V vibratome (Microm, France). Sections encompassing VTA were placed in an ice-cold cutting solution (KCl 2.5 mM, NaH_2_PO_4_ 1.25 mM, MgSO_4_ 10 mM, CaCl_2_ 0.5 mM, glucose 11 mM, sucrose 234 mM, NaHCO_3_ 26 mM) bubbled with 95% O_2_/5% CO_2_. Slices were then incubated in aCSF at 37°C for 1 h, and then kept at RT. For recordings, sections were transferred in a thermo-controlled (32-34°C) recording chamber superfused with aCSF (2.5 ml/min). Spontaneous excitatory post-synaptic currents (sEPSCs) or excitability were measured using visualized whole-cell voltage-clamp and current-clamp recordings, respectively, using an upright microscope (Olympus France). Putative dopamine neurons were identified as previously described [73]. Current-clamp recordings were performed using a Multiclamp 700B device (Molecular Devices, Sunnyvale, CA). Signals were collected and stored using a Digidata 1440A converter and pCLAMP 10.2 software (Molecular Devices, CA). In all cases, access resistance was monitored by a step at −10 mV (0.1 Hz) and experiments were discarded if the access resistance increased more than 20% (internal solution: K-D-gluconate 135 mM, NaCl 5 mM, MgCl_2_ 2 mM, HEPES 10 mM, EGTA 0.5 mM, MgATP 2 mM, NaGTP 0.4 mM). Depolarizing (0-300 pA) or hyperpolarizing (0-450 pA) 800 ms current steps were used to assess excitability and membrane properties of VTA dopamine neurons. sEPSCs were assessed in voltage-clamp mode at a voltage of −65 mV in the presence of picrotoxin (50 μM) using the same internal solution. Off-line analyzes were performed using Clampfit 10.2 (Axon Instruments, USA).

### Fecal microbiota transfer (FMT)

The FMT protocol was adapted from [74]. For induction experiments (FMT*^p^*^-Cresol^ vs. FMT^Control^), recipient C57BL/6J mice were received at 3 weeks of age and habituated for 5 days to our facility. From Day 1 to 3, mice were gavaged with omeprazole (Esomeprazole, Biogaran, 50 mg/Kg/24 h) each morning to lower gastric acidity and favor survival of the inoculum. In the evening of Day 3, food was removed, but mice had *ad libitum* access to water. On Day 4, mice received 5 consecutive gavages of 200 µL of a laxative, reconstituted as recommended by the manufacturer (Moviprep, Norgine SAS, Rueil-Malmaison, France) at 1 hr 30 min intervals. The FMT inoculum was prepared by pooling individual fecal pellets from n=15 control or n=15 mice treated with *p*-Cresol for 4 weeks (as described in the animal treatment section). Fecal pools were then weighted and homogenized in ice-cold ddH_2_O (weight:volume=1:50). Fecal slurry was filtered over a 70 µm mesh to remove fibers and clogs. Recipient mice received 3 consecutive gavages of 200 µL of the filtered fecal slurry at 2 h intervals. After the last gavage, mice were returned to clean cages and had *ad libitum* access to food and water. Body weight was monitored on days 4, 5 and 7 to verify that the mice recovered from the FMT procedure. Three weeks post FMT, mice were subjected to behavioral tests two days apart in the following order: dyadic social interactions, motor stereotypies, novelty-suppressed feeding test and zero-maze.

For rescue experiments (FMT^Control^), mice were treated with *p*-Cresol for 4 wks (as described in the animal treatment section), subjected to behavioral tests two days apart in the following order: direct social interactions, motor stereotypies and three-chamber test. *p*-Cresol treatment was stopped at the end of behavioral testing when Control and *p*-Cresol-treated mice were subjected to the FMT procedure described above. The FMT inoculum was prepared by pooling individual fecal pellets from n=15 control donor untreated male mice of the same age. Three weeks post FMT, mice were subjected to behavioral tests two days apart in the following order: direct social interactions, motor stereotypies, three-chamber test. The *ex vivo* electrophysiological recordings were also performed three weeks post FMT on parallel sub-groups of mice which were not subjected to behavioral testing.

### Fecal microbiota analysis using 16S rRNA sequencing

#### Fecal DNA sample preparation

Mice were individually placed for 10 min in a sterile empty plastic cage, and feces were collected in Eppendorf tubes in the morning (10-11 a.m.), snapped-frozen in liquid nitrogen and stored at −80°C until further use. Genomic DNA was obtained from fecal samples using the QIAamp power fecal DNA kit (Qiagen), and DNA concentration was determined using a TECAN Fluorometer (Qubit® dsDNA HS Assay Kit, Molecular Probes).

#### 16S sequencing

The V3-V4 hypervariable region of the 16S rRNA gene was amplified by PCR using the following primers: a forward primer 5′-**CTT TCC CTA CAC GAC GCT CTT CCG ATC T**AC GGR AGG CAG CAG-3 (28-nt Illumina adapter (in bold) followed the 14 nt broad range bacterial primer 343F) and a reverse primer 5′-**GGA GTT CAG ACG TGT GCT CTT CCG ATC T**TA CCA GGG TAT CTA ATC CT-3′ (28-nt Illumina adapter (in bold) followed by the 19 nt broad range bacterial primer 784R). The PCR reaction mix consisted of 10 ng of fecal DNA template, 0.5 µM primers, 0.2 mM dNTP, and 0.5 U of the DNA-free Taq-polymerase (MolTaq 16S, Molzym, Bremen, Germany). The PCR cycles were as follow: 1 cycle at 94°C for 60 s, followed by 30 cycles at 94°C for 60 s, 65°C for 60 s, 72°C for 60 s, and a final step at 72°C for 10 min. The PCR reactions were sent to the @Bridge platform (INRAe, Jouy-en-Josas, france) for sequencing (Illumina Miseq technology). Single multiplexing was performed using home-made 6 bp index, which were added to R784 during a second PCR with 12 cycles using forward primer (AAT GAT ACG GCG ACC ACC GAG ATC TAC ACT CTT TCC CTA CAC GAC) and reverse primer (CAA GCA GAA GAC GGC ATA CGA GAT-index-GTG ACT GGA GTT CAG ACG TGT). The resulting PCR products were purified and loaded onto the Illumina MiSeq cartridge according to the manufacturer instructions. Run quality was checked internally using PhiX, and sequences assigned to the corresponding based on the 6 pb index. *Sequences preprocessing.* Demultiplexed reads were imported as a whole dataset into QIIME2 [75] (v. 2020.8) that was used for further processing [76]. Primer sequences were removed from reads with Cutadapt [77] and primer-deprived reads were discarded. Based on a sequencing quality score above 18 as recommended [78], forward and reverse reads were truncated at base position 236 and 225 respectively. Paired reads were then processed with QIIME2 implementation of DADA2 [79]. Low quality reads (with more than 2 expected errors) were dropped. Reads were then denoised by the pseudo-pooling method for finer low-count variant detection. Chimera reads were removed using the pooled option. 53% of total reads passed quality control checks and were included in downstream analysis, yielding 2494 Amplicon Sequence Variants (ASV) of which 873 were detected at least in 2 individuals from each condition.

#### Richness and diversity

High-quality reads were aligned to QIIME2 reference library using mafft [80, 81]. Aligned reads were masked in order to remove high variation reads. Then a phylogenetic tree was constructed from the masked alignment of all ASV sequences with the QIIME2 implementation of FastTree [82]. Richness and evenness metrics were calculated with the QIIME2 “richness and diversity” plugin and Observed richness’, Shannon’s and Pielou evenness’ indexes were computed for each condition. Divergence in community composition between samples was quantitatively assessed through a compositional β-diversity metric rooted in a centered log-ratio transformation and matrix completion called robust Aitchison PCA and implemented in QIIME2, which has superior performances to more classical abundance-based Bray-Curtis or Unifrac distances [83]. Aitchison’s distances were calculated on non-rarefied compositional datasets using DEICODE based on 10,453 reads/sample, the largest sampling depth possible, that is robust with respect to compositional data with high levels of sparsity [83]. Hypothesis testing with the beta-group-significance was performed using PERMANOVA applied to the β-diversity ordination artefact using the native QIIME2 implementation of ADONIS from the vegan R package with 999 permutations. Sequences, ASV table and rooted phylogenetic tree were extracted from QIIME2 for further analysis.

#### Taxonomic assignment

To optimize taxonomic affiliation of the ASV, we used a three-step approach. First, taxonomy was inferred using Kraken2 algorithm that is more accurate and faster for 16S rRNA profiling data than the sklearn-classifier implemented in QIIME 2 [84]. Kraken 2 internal quality score was set at 0.05 as recommended [84]. To ascertain first-pass taxonomic affiliation, individual ASV sequences were aligned against the 16S rRNA sequences of their respective Kraken 2-derived taxonomic affiliation using NCBI BLAST+ from the non-recombinant nucleotide (nr/nt) database architecture [85]. 75% of ASV yielded first-pass satisfactory taxonomic assignment with> 95% homology with their first-hit assigned sequence. The remaining 25% of ASV were blasted again against the next hit proposed by Kraken2 until a higher homology score was reached in a method inspired by [86]. After these iterative steps, we assigned up to 85% of all ASV with a homology score above 95%. For the remaining unassigned ASV (15%), their sequence was blasted on the nr/nt database of bacterial sequences excluding uncultured and environmental sequences, yielding taxonomic assignation for 7% ASV more. At the end of the assignment process, 92% of ASV were taxonomically assigned with a homology score above 95%. For the remaining 8% of ASV with no hit above 95% homology, the closest homology match sequence (including uncultured and environmental sequences) was recorded. Tab. S4 recapitulates the name and sequence of each of the ASVs detected in at least two individuals in each dataset, as well as the taxon identity and homology scores (% of identity, bitscore, e-value) for the nearest homology match. *Microbial composition analysis.* Individual datasets management (taxonomy, abundance and metadata) was done on Phyloseq [87]. The effects of *p*-Cresol treatment or FMT*^p^*^-Cresol^ vs. FMT^Control^ were assessed using the Analysis of Composition of Microbiomes (ANCOM) method [88]. ANCOM accounts for the compositional nature of 16S datasets, is very sensitive for datasets >20 samples and displays superior performances to control for false discovery rate compared to other methods such as DESeq2 [37]. The utility of ANCOM has been demonstrated for preprocessing sparse microbiome data sets with matrix completion to allow compositional ordination and to preserve information about the features driving differences among samples [37]. ANCOM accounts for compositionality using centered log-ratio (CLR) analysis and therefore improves inference in microbiota survey data [37]. Recommended ANCOM thresholds were used to identify dysregulated taxa: |CLR|>0.2 and W=0.7. Of note, the *p*-Cresol versus Control dataset consisted of two batches of sequences obtained from two independent biological replicates (n=15 animals/group/replicate) and the built-in batch effect correction implemented in ANCOM was used to account for batch effect. Based on features identified by ANCOM at each taxonomic level (from ASV to phylum), synthetic cladograms were built using GraPhlAn [89].

#### Association between microbiota composition, behavioral scores or fecal p-Cresol

To assess relationships between changes in microbial distributions and behavioral variables or fecal *p*-Cresol), we used Random Forest regression models with nested cross-validation implemented in QIIME2. Independent models were built to identify ASV best predicting social interaction deficits, using the variable “total social contact time” or stereotypies using a composite variable obtained by summing the centered-scaled numbers of circling events and headshakes or fecal *p*-Cresol concentrations.

#### Blast analysis of p-Cresol biosynthetic enzymes

As references for queries, we used the protein sequence of ThiH (tyrosine lyase (NP 418417) from *E. coli* K-12 substrate MG1655), TyrB (tyrosine aminotransferase (NP 418478) from *E. coli* K-12 substrate MG1655) and HpdA, HpdB and HpdC (*p*-hydroxyphenylacetate decarboxylase subunits A (AJ543427), B (AJ543425) and C (AJ543426) from *C. difficile* DSM 1296T) (Tab. S3). Sequence homology search was performed using the Blast webtool set with default parameters on the nr/nt database restricted to *Duncaniella dubosii* (taxid:2518971) and the genera *Barnesiella* (taxid:397864) and *Anaerobium* (taxid:1855714). We considered as potential homologous enzymes sequences presenting amino acid identity above 30% along more than 50% of the query sequence.

### Statistics

Comparisons between two groups were performed using 2-tailed unpaired Mann-Whitney’s non-parametric U-test. Multiple group comparisons were performed using two-way ANOVA with factors stated in legends and text. Prior to ANOVA, normality of data was assessed using Kolmogorov-Smirnov’s test. *Post hoc* comparisons were performed using Šidák’s correction for multiple comparison. Detailed ANOVA statistics for each panel are provided in Tab. S5. For comparison analysis of frequency distributions, Kolmogorov-Smirnov’s test was used. Principal Component Analysis (PCA) of behavioral data was performed for each dataset consisting of scores from the dyadic social interaction test and scores from the direct monitoring of motor stereotypies using the webtool Clustvis [90]. Statistical significance was set according to a two-tailed p-value or adjusted (if relevant) p-value(p)<0.05. Only significant differences are displayed on the graphs.

## Consent for publication

Not applicable.

## Availability of data and material

The datasets acquired and analyzed during the current study are available from the corresponding author on reasonable request.

## Authors’ contributions

PBM, JAJB, JCa, JCh and JLM performed behavioral phenotyping and scoring. SPF, RCC and JB performed electrophysiological recordings. BA, CC and PL performed 16S rRNA sequencing. NC and SB performed the analysis of 16S rRNA datasets. LMG, AM and MED performed urinary and serum *p*-Cresol measures. JML and JC performed fecal *p*-Cresol measures. LD designed the study, performed statistical analysis and wrote the manuscript. NG contributed to study design and writing of the manuscript. All authors read and approved the final manuscript.

## Acknowledgements

We thank Lucien Relmy for expert technical assistance with animal care and Thomas Lorivel for help with behavioral tests set-up. We are grateful to Dominique Gauguier (Centre de Recherche des Cordeliers, Paris) for helpful discussions regarding the procedure for fecal microbiota transfers.

## TABLES

**Tab. 1.** Predicted proteins homologous to enzymes involved in metabolic pathways from tyrosine to *p*-Cresol in species related to bacterial taxa identified as upregulated both upon *p*-Cresol treatment and upon FMT*^p^*^-Cresol^. Reference enzyme protein sequences from ThiH (NP_418417.1, 2-iminoacetate synthase from *Escherichia coli* K12, metabolizing tyrosine to *p*-Cresol) and HpdA (sp|Q84F14|HPDA_CLODI, 4-hydroxyphenylacetate decarboxylase activating enzyme from *Clostridioides difficile*, metabolizing 4-hydroxyphenylacteate to *p*-Cresol) were used to conduct a Blast search against sequences from the *Anaerobium* genus and from the *Muribaculaceae* family. Only sequences covering more than 50% of the query sequence and presenting amino-acid identity above 30% were considered as potential homologous enzymes. See also Tab. S3 for the full blast analysis.

| Species | Direct pathway |  |  | Indirect pathway |  |  |
| --- | --- | --- | --- | --- | --- | --- |
|  | ThiH |  |  | HpdA |  |  |
|  | Accession | % Identity | E-score | Accession | % Identity | E-score |
| *Anaerobium acetethylicum* | WP_091230597.1 | 38.90% | 3.00E-72 | WP_091236185.1 | 37.62% | 4E-66 |
| *Duncaniella dubosii* | WP_123615573.1 | 46.85% | 3.00E-121 | WP_123485485.1 | 32.59% | 5E-13 |
| *Barnesiella* sp. | MBD5257534.1 | 48.49% | 1.00E-129 | - | - | - |
| *Barnesiella* sp. An55 | WP_087424377.1 | 49.04% | 3.00E-125 | - | - | - |
| *Barnesiella intestinhominis* | WP_195342987.1 | 48.65% | 2.00E-124 | - | - | - |
| *Barnesiella viscericola* | WP_025277286.1 | 47.95% | 6.00E-123 | - | - | - |

## SUPPLEMENTARY FIGURES

**Fig. S1.**
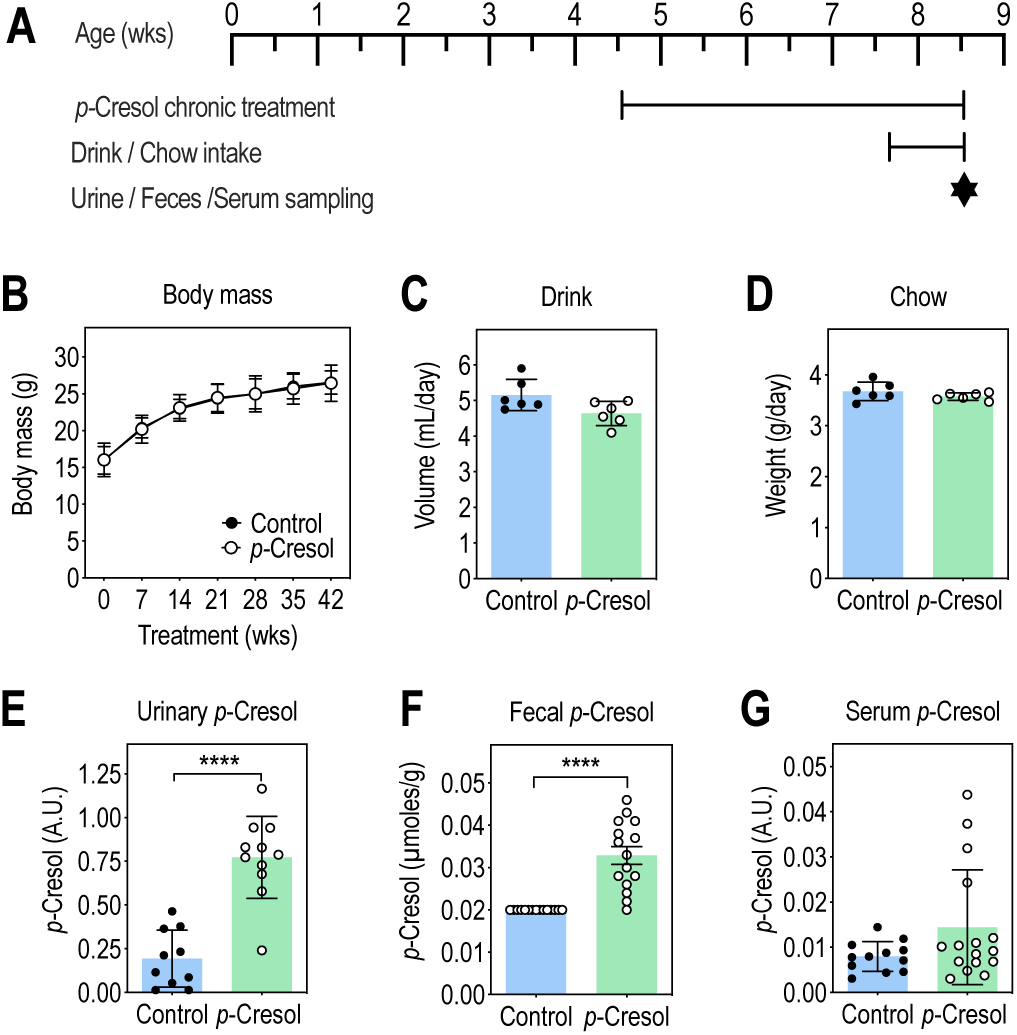
*p-*Cresol treatment does not impact general physiological parameters, increases urinary and fecal *p-* Cresol excretion but not its serum levels. (A) Timeline of the experiment (B) Body weight follow-up (n=24/group); 2-way ANOVA: p(Treatment)=0.8884, p(Time)<0.0001, p(Treatment x Time)=0.9971; Šidák’s post hoc tests for treatment effect: p>0.99. (C) Drink intake (n=6/group, each point representing the mean consumption per animal in one cage of 5 animals); Mann-Whitney U-test: p=0.0931. (D) Food intake (n=6/group, each point representing the mean consumption per animal in one cage of 5 animals); Mann-Whitney U-test: p=0.3939. (E) Urinary levels of *p*-Cresol after 4 weeks treatment (n=10 Control, n=11 *p-*Cresol); Mann-Whitney U-test: ****, p<0.0001. (F) Fecal levels of *p*-Cresol after 4 weeks treatment (n=15 Control, n=15 *p-*Cresol); Mann-Whitney U-test: ****, p<0.0001. (G) Serum levels of *p*-Cresol after 4 weeks treatment (n=12 Control, n=16 *p-*Cresol); Mann-Whitney U-test: p=0.3238. (B) Data are presented as means ± SD. (C-G) Data are presented as dot-plots featuring means ± SD.

**Fig. S2.**
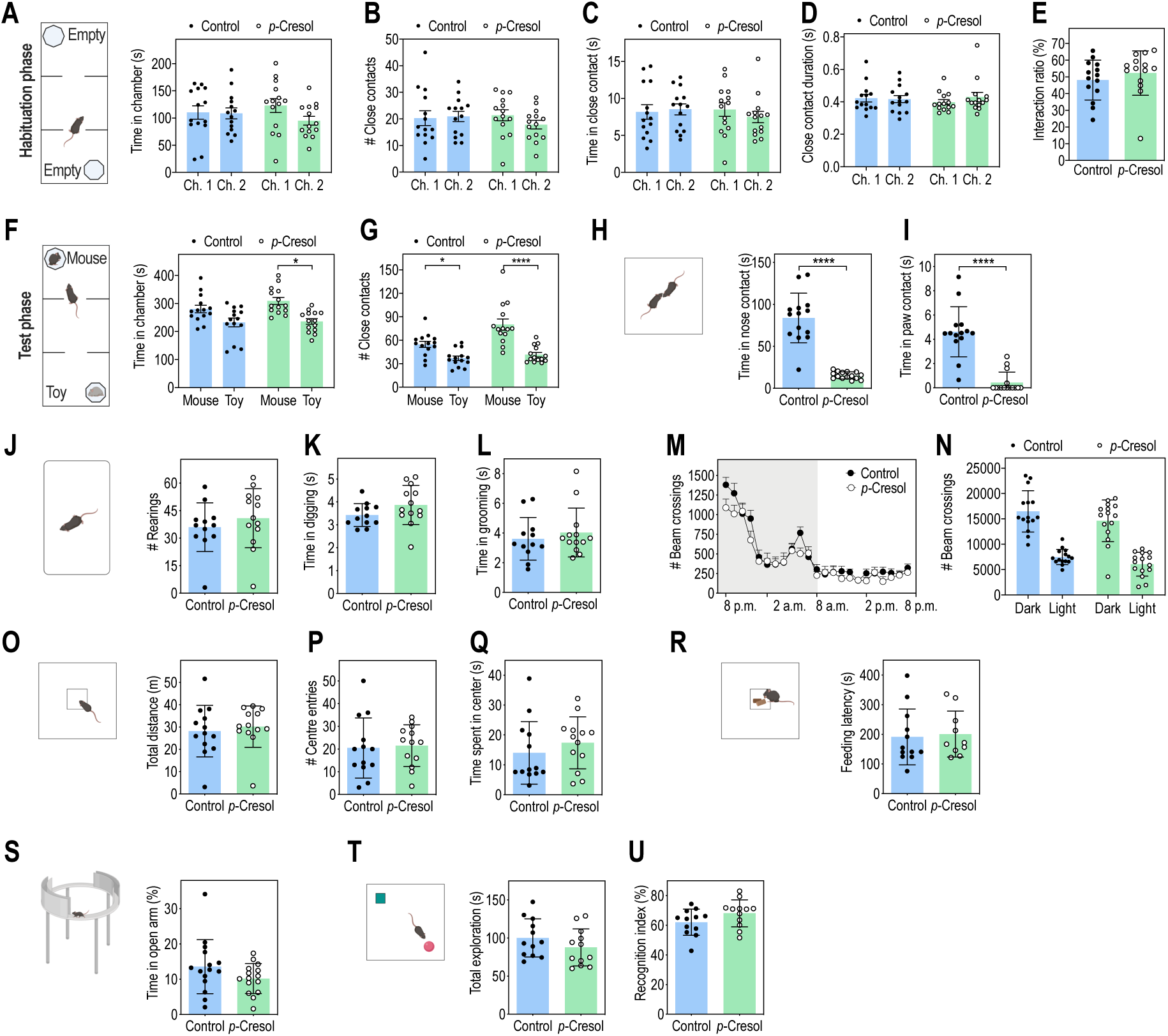
*p*-Cresol treatment-induced behavioral alterations: additional parameters in the 3-chamber, dyadic social interactions and stereotypies tests and results from tests assessing locomotor activity, anxiety or cognition (relative to Fig. 1) (A-E) Additional parameters during the habituation phase in the 3-chamber test (n=14/group): (A) Time spent in each chamber; 2-way ANOVA: p(Treatment)=0.9498, p(Chamber)=0.3018, p(Treatment x Chamber)=0.3599; Šidák’s post hoc tests for chamber effect: p>0.05. (B) Number of close contacts to the empty wired cage in each chamber; 2-way ANOVA: p(Treatment)=0.6839, p(Chamber)=0.4390, p(Treatment x Chamber)=0.2648; Šidák’s post hoc tests for chamber effect: p>0.05. (C) Time close contacts to the empty box in each chamber; 2-way ANOVA: p(Treatment)=0.7265, p(Chamber)=0.6260, p(Treatment x Chamber)=0.3188; Šidák’s post hoc tests for chamber effect: p>0.05. (D) Mean duration of each close contacts to the empty box in each chamber; 2-way ANOVA: p(Treatment)=0.8330, p(Chamber)=0.4995, p(Treatment x Chamber)=0.3378; Šidák’s post hoc tests for chamber effect: p>0.05. (E) Interaction ratio; Mann-Whitney U-test: p=0.265. (F, G) Additional parameters during the test phase in the 3-chamber test: (F) Time spent in each chamber containing the mouse interactor or the toy mouse; 2-way ANOVA: p(Treatment)=0.0061, p(Mouse-Toy)=0.0019, p(Treatment x Mouse-Toy)=0.4797; Šidák’s post hoc tests for Mouse vs. Toy preference effect: p>0.05 for Control group, * p<0.05 for *p*-Cresol group. (G) Number of close contacts with the mouse interactor or the toy mouse; 2-way ANOVA: p(Treatment)=0.0035, p(Mouse-Toy)<0.0001, p(Treatment x Mouse-Toy)=0.0257; Šidák’s post hoc tests for Mouse vs. Toy preference effect: *p<0.05, ****p<0.0001. (H, I) Additional parameters in the dyadic social interaction test (n = 14 Control, n=15 *p*-Cresol): (H) Total time spent in nose and (I) paw contact; Mann-Whitney U-test: ****p<0.0001. (J-L) Additional parameters in the motor stereotypies test (n=11 Control, n=12 *p*-Cresol): (J) Number of rearings; Mann-Whitney U-test: p=0.3400. (K) Time spent in digging; Mann-Whitney U-test: p=0.4613. (L) Time spent in grooming; Mann-Whitney U-test: p=0.1366. (M, N) Locomotor activity in actimetry chambers (n=15/group): (M) Horizontal activity follow-up over 24 h; 2-way ANOVA: p(Treatment)=0.1057, p(Time)<0.0001, p(Treatment x Phase)=0.3610; Šidák’s post hoc tests for treatment effect: p>0.05. (N) Cumulative activity in the dark or light phase; 2-way ANOVA: p(Treatment)=0.0909, p(Phase)<0.0001, p(Treatment x Phase)=0.7513; Šidák’s post hoc tests for treatment effect: p>0.05. (O-Q) Open-field test (n=13/group); (O) Total distance travelled; Mann-Whitney U-test: p=0.3107. (P) Number of center entries; Mann-Whitney U-test: p=0.6047. (Q) Time spent in the center of the area; Mann-Whitney U-test: p=0.3292. (R) Novelty suppressed feeding (NSF) test (n=12 Control, n=10 *p*-Cresol): latency to first bite; Mann-Whitney U-test: p=0.5707 (S) Zero-maze test (n=15 Control, n=15 *p*-Cresol): percentage of time spent in open arm; Mann-Whitney U-test: p=0.1485. (T, U) Novel object recognition (NOR) (n=12/group): (T) Total time spent exploring the objects during the test phase; Mann-Whitney U-test: p=0.1978. (U) % of time spent exploring the old vs. the new object; Mann-Whitney U-test: p=0.1135. (A-L, N-U) Data are presented as dot-plots featuring means ± SD. (M) Data are shown as means ± SD.

**Fig. S3.**
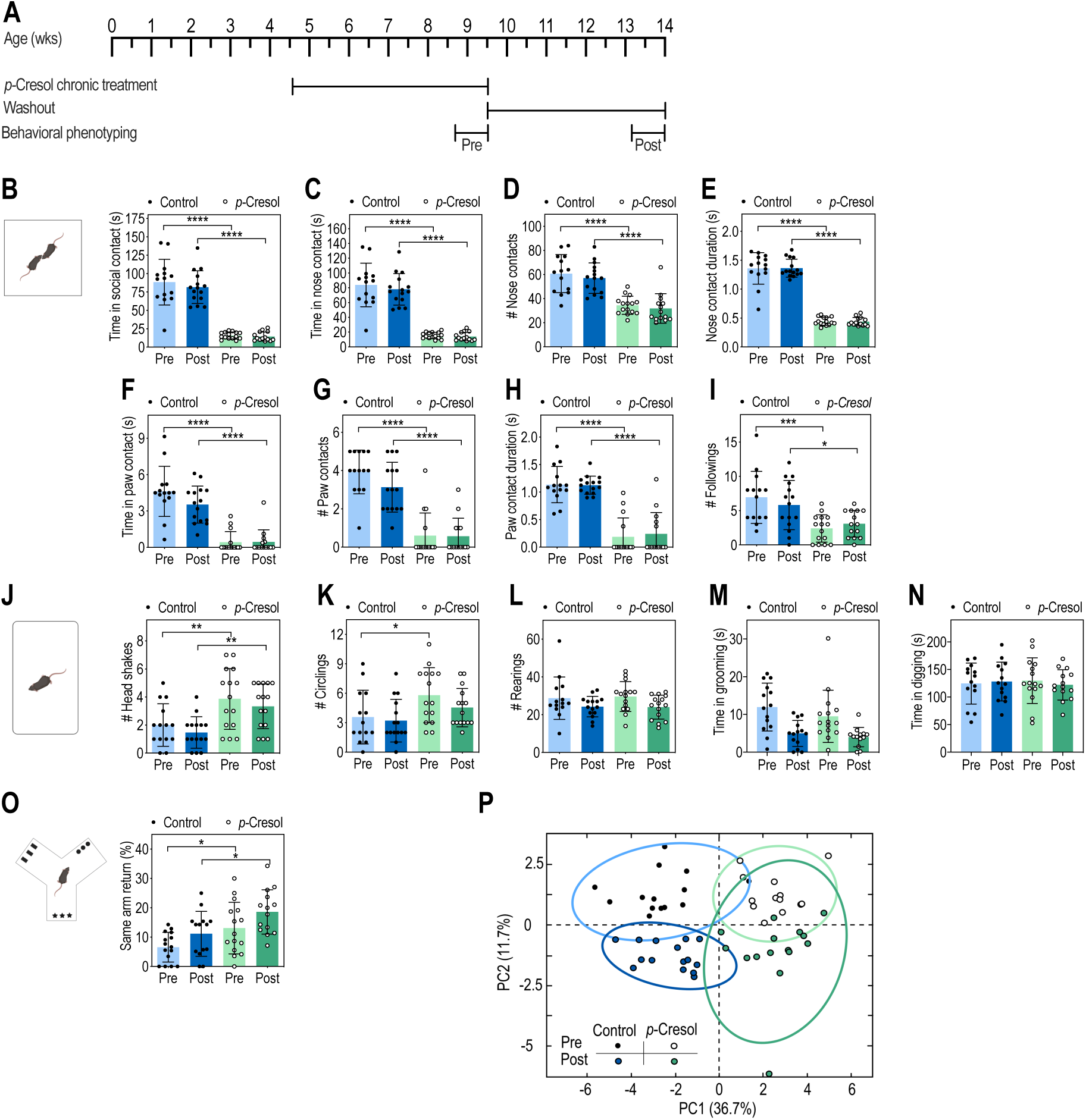
Autistic-like behaviors persist after discontinuation of *p*-Cresol for 4 weeks. (A) Timeline of the experiment (B-G) Dyadic social interaction test (n=14 Control, n=15 *p*-Cresol): (B) Total time spent in social contact; 2-way ANOVA: p(Treatment)<0.0001, p(Washout)=0.4194, p(Treatment x Washout)=0.5883; Šidák’s post hoc tests for treatment effect: ****p<0.0001. (C) Time spent in nose contact; 2-way ANOVA: p(Treatment)<0.0001, p(Washout)=0.4525, p(Treatment x Washout)=0.6450; Šidák’s post hoc tests for treatment effect: ****p<0.0001. (D) Number of nose contacts; 2-way ANOVA: p(Treatment)<0.0001, p(Washout)=0.3237, p(Treatment x Washout)=0.8660; Šidák’s post hoc tests for treatment effect: ****p<0.0001. (E) Mean duration of each nose contact; 2-way ANOVA: p(Treatment)<0.0001, p(Washout)=0.9807, p(Treatment x Washout)=0.8922; Šidák’s post hoc tests for treatment effect: ****p<0.0001. (F) Time spent in paw contact; 2-way ANOVA: p(Treatment)<0.0001, p(Washout)=0.1638, p(Treatment x Washout)=0.1465; Šidák’s post hoc tests for treatment effect: ****p<0.0001. (G) Number of paw contacts; 2-way ANOVA: p(Treatment)<0.0001, p(Washout)=0.1793, p(Treatment x Washout)=0.2109; Šidák’s post hoc tests for treatment effect: ****p<0.0001. (H) Mean duration of each paw contact; 2-way ANOVA: p(Treatment)<0.0001, p(Washout)=0.7906, p(Treatment x Washout)=0.6926; Šidák’s post hoc tests for treatment effect: ****p<0.0001. (I) Number of followings, 2-way ANOVA: p(Treatment)<0.0001, p(Washout)=0.7697, p(Treatment x Washout)=0.2517; Šidák’s post hoc tests for treatment effect: ***p<0.001, *p<0.05. (J-M) Motor stereotypies (n=14 Control; n=15 *p-*Cresol): (J) Number of head shakes; 2-way ANOVA: p(Treatment)<0.0001, p(Washout)=0.2185, p(Treatment x Washout)>0.9999; Šidák’s post hoc tests for treatment effect: **p<0.01. (K) Number of circling events; 2-way ANOVA: p(Treatment)=0.0051, p(Washout)=0.2605, p(Treatment x Washout)=0.5796; Šidák’s post hoc tests for treatment effect: *p<0.05 for Control, p>0.05 for *p-*Cresol. (L) Number of rearings; 2-way ANOVA: p(Treatment)=0.8820, p(Washout)=0.0188, p(Treatment x Washout)=0.7823; Šidák’s post hoc tests for treatment effect: p>0.05. (M) Time spent in grooming; 2-way ANOVA: p(Treatment)=0.2308, p(Washout)<0.0001, p(Treatment x Washout)=0.5929; Šidák’s post hoc tests for treatment effect: p>0.05. (N) Time spent in digging; 2-way ANOVA: p(Treatment)=0.9763, p(Washout) = 0.8304, p(Treatment x Washout)= 0.5579; Šidák’s post hoc tests for treatment effect: p>0.05. (N) Y-maze spontaneous alternations: percentage of same arm returns (n=15 Control, n=14 *p*-Cresol): 2-way ANOVA: p(Treatment) = 0.0007, p(Washout) = 0.0120, p(Interaction) = 0.8179; Šidák’s post hoc tests for treatment effect: *p<0.05. (P) PCA plots of behavioral scores recorded in the dyadic social interaction and direct monitoring of motor stereotypies (Fig. S3B-N); ellipses of the 95% confidence intervals are indicated for each group (n=14 Control; n=15 *p-*Cresol). (B-O) Data are presented as dot-plots featuring means ± SD.

**Fig. S4.**
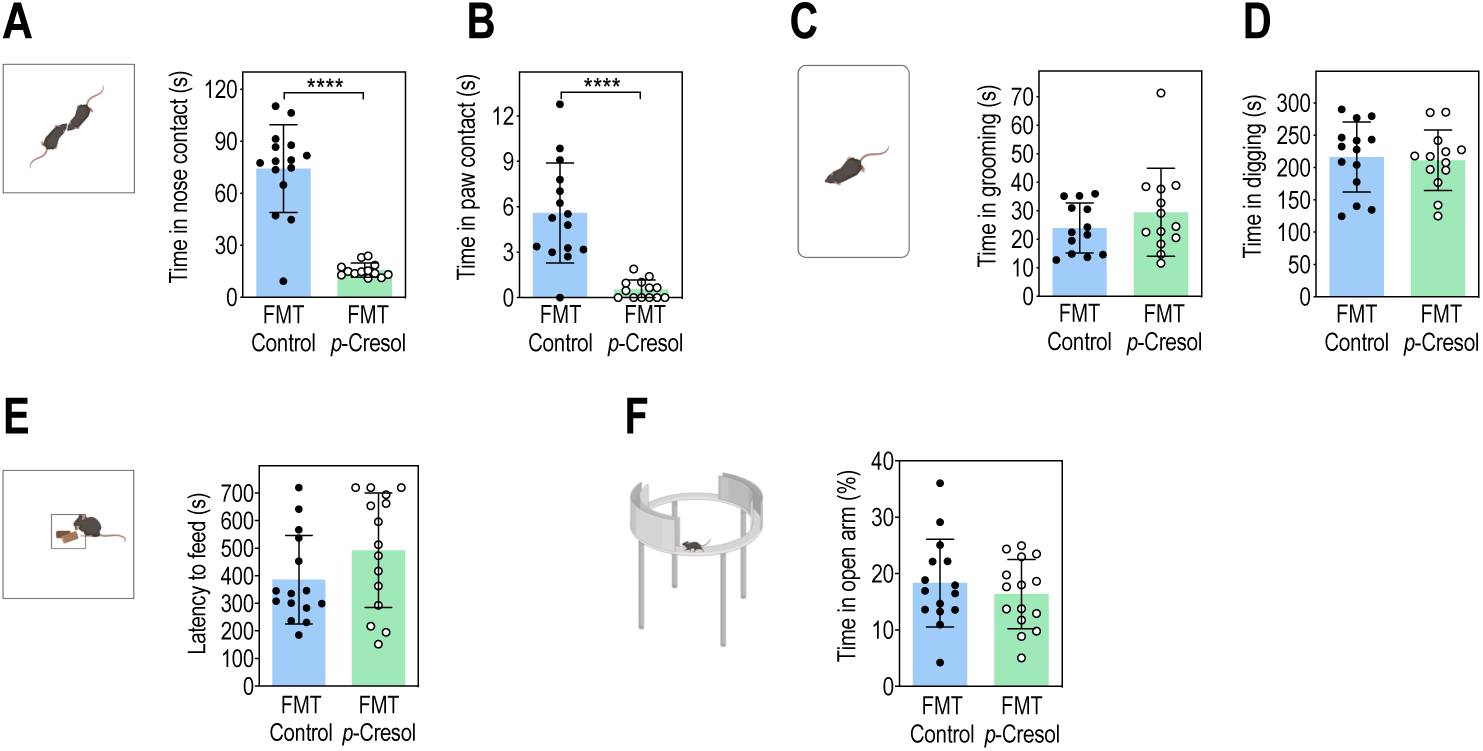
Additional behavioral parameters for dyadic social interactions, stereotypies tests and anxiety in FMT^Control^ and FMT*^p-^*^Cresol^ mice (relative to Fig. 3) (A, B) Dyadic social interaction test (n=15 animals/group): (A) Total time spent in nose contact; Mann-Whitney U-test: ****p<0.0001 (B) Total time spent in paw contact; Mann-Whitney U-test: ****p<0.0001 (C, D) Motor stereotypies (n= 14 FMT^Control^, n=13 FMT*^p^*^-Cresol^): (C) Time spent in grooming; Mann-Whitney U-test: p=0.3358. (D) Time spent in digging; Mann-Whitney U-test: p=0.5502. (E) Novelty suppressed feeding (NSF) test (n=15 FMT^Control^, n=15 FMT*^p^*^-Cresol^): latency to first bite; Mann-Whitney U-test: p=0.164. (F) Zero-maze test (n=15 FMT^Control^, n=15 FMT*^p^*^-Cresol^): percentage of time spent in open arm; Mann-Whitney U-test: p=0.653. Data are presented as dot-plots featuring means ± SD.

**Fig. S5.**
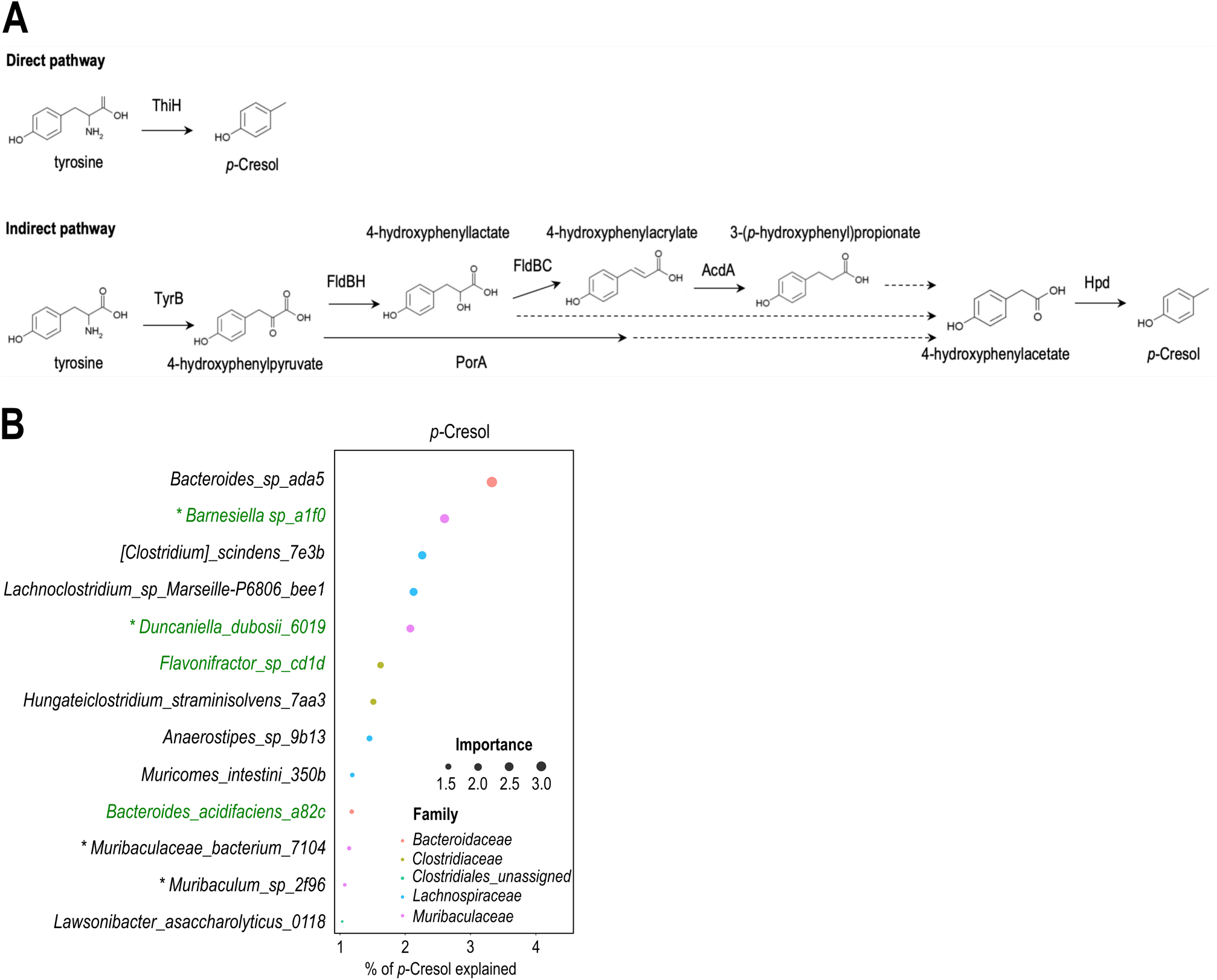
*p-*Cresol microbial biosynthetic pathways (relative to Fig. 4) (A) Metabolic pathways from tyrosine to *p*-Cresol (C, D). The direct pathway uses thiazole synthase H (ThiH), while the indirect pathway uses several intermediate enzymes: tyrosine aminotransferase B (TyrB), phenyllactate dehydrogenase (FldH), phenyllactate dehydratase (FldBC), acyl-CoA dehydrogenase (AcdA), hydroxyarylic acid decarboxylase (Had), pyruvate-ferredoxin oxidoreductase A (PorA) and hydroxyphenylacetate decarboxylase (Hpd). Dashed lines indicate enzymes not yet identified/characterized. Adapted from [28]. (B) Microbial composition prediction of fecal *p*-Cresol in FMT^Control^ and FMT*^p-^*^Cresol^ mice (n=8 FMT^Control^, n=12 FMT*^p-^*^Cresol^). ASV best predicting fecal *p*-Cresol as identified by Random Forest analysis. Only ASV contributing >1% accuracy in fecal *p*-Cresol concentration prediction are presented. ASV related to ASV/species increased in FMT*^p^*^-Cresol^ mice as determined by ANCOM analysis are labeled in green. Asterisks indicate members of the *Muribaculaceae* family.

**Fig. S6.**
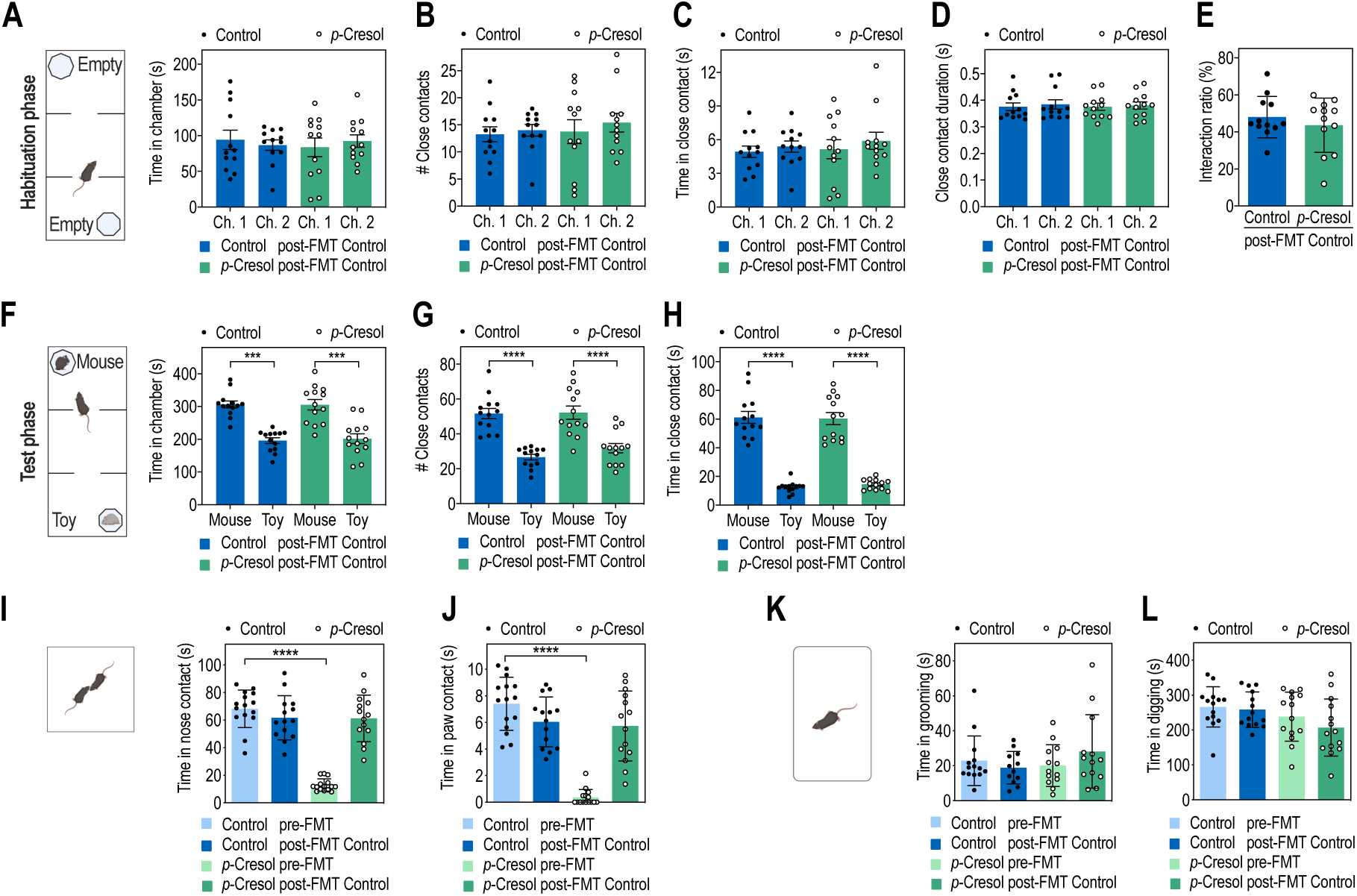
FMT experiments from control donors (FMT^Control^) to Control or *p*-Cresol-treated recipient mice: additional behavioral parameters in the three-chamber, dyadic social interactions and stereotypies tests (relative to Fig. 5) (A-E) Additional parameters during the habituation phase in the 3-chamber test post-FMT^Control^ (n=13/group): (A) Time spent in each chamber; 2-way ANOVA: p(Treatment)=0.8296, p(Chamber)=0.9514, p(Treatment x Chamber)=0.4749; Šidák’s post hoc tests for chamber effect: p>0.05. (B) Number of close contacts with the empty box in each chamber; 2-way ANOVA: p(Treatment)=0.6456, p(Chamber)=0.2947, p(Treatment x Chamber)=0.6878; Šidák’s post hoc tests for chamber effect: p>0.05. (C) Time in close contact with the empty box in each chamber; 2-way ANOVA: p(Treatment)=0.6728, p(Chamber)=0.1662, p(Treatment x Chamber)=0.7417; Šidák’s post hoc tests for chamber effect: p>0.05. (D) Mean duration of each close contact to the empty box in each chamber; 2-way ANOVA: p(Treatment)=0.9252, p(Chamber)=0.2163, p(Treatment x Chamber)=0.7061; Šidák’s post hoc tests for chamber effect: p>0.05. (E) Interaction ratio for Chamber 1 over Chamber 2; Mann-Whitney U-test: p=0.755. (F-H) Additional parameters during the test phase in the 3-chamber test post-FMT^Control^ (n=13/group) (F) Time spent in each chamber containing the mouse interactor or the toy mouse; 2-way ANOVA: p(Treatment)=0.7588, p(Mouse-Toy) <0.0001, p(Treatment x Mouse-Toy)=0.8356; Šidák’s post hoc tests for Mouse vs. Toy preference: ***p<0.001. (G) Number of close contacts with the mouse interactor or the toy mouse; 2-way ANOVA: p(Treatment)=0.3399, p(Mouse-Toy)<0.0001, p(Treatment x Mouse-Toy)=0.4225; Šidák’s post hoc tests for Mouse vs. Toy preference: ****p<0.0001. (H) Time in close contact with the mouse interactor or the toy mouse; 2-way ANOVA: p(Treatment)=0.8480, p(Mouse-Toy)<0.0001, p(Treatment x Mouse-Toy)=0.6498; Šidák’s post hoc tests for Mouse vs. Toy preference: ****p<0.0001. (I, J) Additional parameters in the dyadic social interaction test pre- and post-FMT^Control^ (n=15/group pre-FMT, n=14/group post-FMT): (I) Time spent in nose contact; 2-way ANOVA: p(Treatment)<0.0001, p(FMT^Control^)<0.0001, p(Treatment x FMT^Control^) <0.0001; Šidák’s post hoc tests for treatment effect: ****p<0.0001 for pre-FMT^Control^ groups, p>0.05 post- FMT^Control^ groups. (J) Time spent in paw contact; 2-way ANOVA: p(Treatment)<0.0001, p(FMT^Control^)=0.0002, p(Treatment x FMT^Control^)<0.0001; Šidák’s post hoc tests for treatment effect: ****p<0.0001 pre-FMT Control, groups p>0.05 post-FMT Control groups. (K, L) Additional parameters in the motor stereotypies test pre- and post-FMT^Control^ (n=14/group pre-FMT, n=14/group post-FMT): (K) Time spent in grooming; 2-way ANOVA: p(Treatment)=0.7605, p(FMT^Control^)=0.2801, p(Treatment x FMT^Control^)=0.9682; Šidák’s post hoc tests for treatment effect: p>0.05. (L) Time spent in digging; 2-way ANOVA: p(Treatment)=0.0295, p(FMT^Control^)=0.274, p(Treatment x FMT^Control^)=0.5058; Šidák’s post hoc tests for treatment effect: p>0.05. Data are presented as dot-plots featuring means ± SD.

## SUPPLEMENTARY TABLES LEGENDS

**Tab. S1. ANCOM analysis: significant microbial features (from ASV to phylum) discriminating *p-*Cresol from Control microbiota (relative to Fig. 2)**

Taxonomic information regarding the identified bacterial features, corresponding centered log ratio (CLR), Wald’s test W statistic as determined by ANCOM analysis of bacterial composition based on 16S rRNA sequencing are indicated. ASV, amplicon sequence variant (n=30 Control, n=30 *p-* Cresol).

*Available as Excel file*

**Tab. S2. ANCOM analysis: significant microbial features (from ASV to phylum) discriminating FMT^Control^ from FMT*^p-^*^Cresol^ mice, 3 weeks post FMT (relative to Fig. 4)**

Taxonomic information regarding the identified bacterial features, corresponding centered log ratio (CLR), Wald’s test W statistic as determined by ANCOM analysis of bacterial composition based on 16S rRNA sequencing are indicated. ASV, amplicon sequence variant (n=15 FMT^Control^, n=15 FMT*^p-^*_Cresol)._

*Available as Excel file*

**Tab. S3. Output of blast sequence analysis for ThiH, Tyr and HpdA/B/C enzymes involved in *p*-Cresol synthesis (relative to Tab. 1)**

*Available as Excel file*

**Tab. S4. Details on ASV sequences and taxonomic affiliation.** Name and sequence of each of the ASV identified in the dataset, as well as the taxon identity (txid) and homology scores (% of identity, bitscore, e-value) for the nearest homology match.

*Available as Excel file*

**Tab. S5. Compilation of ANOVA statistics for behavioral data analyses and PERMANOVA statistics for 16S richness and diversity analyses**

*Available as Excel file*

